# Cell signaling pathways discovery from multi-modal data

**DOI:** 10.1101/2025.02.06.636961

**Authors:** Changhan He, Claire Simpson, Ian Cossentino, Bin Zhang, Sasha Tkachev, Devon J. Eddins, Astrid Kosters, Junkai Yang, Shivani Sheth, Tyler Levy, Anthony Possemato, Linglin Huang, Evgeniy Tabatsky, Soo Hee Lee, Debopam Ghosh, Ashley F. George, Ivan Gregoretti, Majd Ariss, Deepti Dandekar, Aniket Ausekar, Nadia R. Roan, Eliver E. B. Ghosn, Marco Colonna, Klarisa Rikova, Qing Nie, Darya Orlova

## Abstract

Deciphering cell signaling pathways is key to understanding biology, disease mechanisms, and developing new therapies. Although advances in multi-omics technologies provide richer insight into signaling, the data remain high-dimensional, heterogeneous, and difficult to interpret, and current computational tools for inferring signaling pathways are limited. To address this, we developed Incytr, a method for efficient discovery of cell signaling pathways through integration of diverse data modalities, including transcriptomics, ATAC-seq, proteomics, phosphoproteomics, and kinomics. We demonstrate its application in COVID-19, Alzheimer’s disease, and cancer, where it successfully recovers known pathways and generates novel, cell-type-specific hypotheses supported by multiple data types. We further show how integrating Incytr-derived pathways with biomarker and drug databases can support target and drug discovery. Finally, we show that using Incytr-derived signaling pathways as training data for simple natural language processing models can deepen our understanding of cell–cell communication and immune cell dynamics, while helping identify new therapeutic targets.

## Introduction

Cell-cell communication via ligand-receptor interactions involves the binding of signaling molecules (ligands) presented or released by sender cells to specific receptors on target cells (or themselves). The specificity of response profiles to ligands is shaped, among other factors, by a combination of selected intracellular signaling intermediates and their concentration distribution, that ultimately leads to activation/inhibition of downstream genes essential for biological processes such as growth, immune response, and tissue repair [Armingol et al., 2020; Wilk et al., 2023]. These signaling pathways involve multiple layers of signal transduction processes, including ligand-receptor interactions, downstream components (e.g., effector molecules) that mediate the cellular response, and target gene activation. Inference and analysis of such pathways is crucial for understanding how cells respond to various stimuli and regulate their functions in both homeostasis and disease [Armingol et al., 2024]. This analysis opens doors to discovering therapeutically relevant pathways [AlMusawi et al., 2021], understanding mechanisms of action [Armingol et al., 2020], and enabling regenerative and personalized medicine approaches [Liu et al., 2021].

For over a decade, RNA-seq has been used as a primary technology for inferring cell signaling pathways, extrapolating key signaling players from gene expression. Thus, the majority of computational methods reported in the literature to date utilize primarily this data modality. A traditional way to infer cell signaling pathways is Gene Set Enrichment Analysis (GSEA), which involves grouping gene expression into signatures, which can then be mapped onto a database of known cell signaling pathways [Subramanian et al., 2005]. While convenient, this method depends on prior knowledge of cell signaling pathways and does not allow *de novo* pathway inference. To facilitate *de novo* signaling pathway inference from gene expression data, methods such as NicheNet [Browaeys et al., 2019], exFINDER [He et al., 2023], and scMLnet [Cheng et al., 2020] that build multi-layer signaling pathways have been developed.

However, recent advancements in newer omics technologies (such as proteomics, and scATAC-seq) enable us to collect more evidence (chromatin accessibility, protein expression, and protein modification) and capture cell signaling information in a more meaningful context than when RNA-seq was the primary source of information. This creates the need for the development of methods that can infer signaling pathways from multi-modal data.

With very few exceptions, current cell signaling pathway inference methods that do not rely on prior knowledge of signaling pathways and that are capable of handling multi-omics data consist of dimensionality reduction and clustering methods using pre-processed multi-modal data. The most recent pre-processing methods involving deep learning approaches (MultiVI [Ashuach et al., 2023], scMoGNN [Wen et al., 2022], scDART [Zhang et al., 2022]) can effectively integrate and leverage information from multiple data modalities simultaneously. These methods focus on constructing joint representations that capture information from multiple modalities (such as scRNA-seq, scATAC-seq, and CITE-seq) for downstream tasks, including clustering and GSEA. However, these approaches do not construct a cell signaling pathway *per se*.

One interesting class of approaches that are independent of prior knowledge of signaling pathways and can be applied to multi-modal data are those based on correlation and stoichiometry scores, such as weighted correlation network analysis (WGCNA) [Langfelder et al., 2008] and de novo multi-omics pathway analysis (DMPA) [Vaparanta et al., 2024]. They integrate and analyze data from multiple modalities, providing a holistic view of cellular processes, and facilitate the construction of co-expression pathways, helping to identify modules of highly correlated genes. However, since correlations do not imply causation, high stoichiometry scores might be misinterpreted as strong associations, even if the underlying biological relevance is weak. One approach to mitigating this issue is to condition gene co-expression on existing or newly generated knowledge of protein-protein interactions (PPIs), including both direct interactions and those that co-occur within the same pathways. This accounts for dependencies in their concentrations and their functional roles, such as ligand-receptor, receptor-effector, or effector-target interactions (Figure 1A). Methods such as NicheNet [Browaeys et al., 2019] and exFINDER [He et al., 2023] have successfully used this strategy, leveraging curated databases of pairwise protein interactions to infer *de novo* intercellular communication by linking ligands to target genes. However, this approach has not yet been applied beyond transcriptomics data.

**Figure 1.**
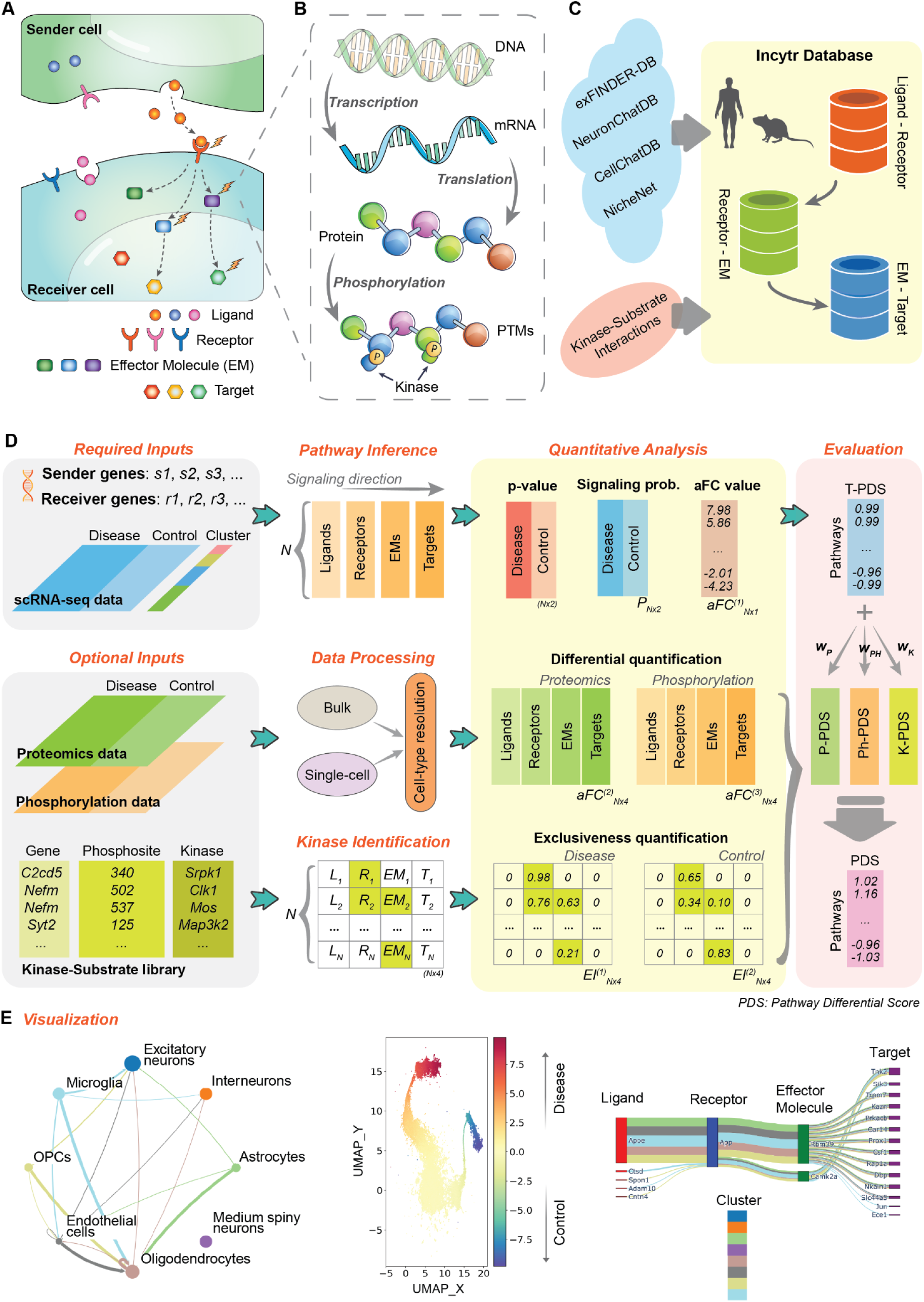
Overview of Incytr. **A.** Cell-cell communication via ligand-receptor binding can involve downstream molecules, forming a ligand-receptor-effector molecule-target (L-R-EM-T) signaling pathway. More generally, this can extend to L–R–…–EM–…–T, as protein–protein interaction databases include both direct and indirect interactions, including those mediated by associations or co-expression within shared pathways. **B.** For an involved molecule to perform its proper functions, multiple steps should be considered, including DNA-RNA transcription, RNA-protein translation, and, in some cases, protein phosphorylation. **C.** The Incytr database (IncytrDB) is integrated from multiple publicly deposited resources and consists of ligand-receptor, receptor-EM, and EM-target interactions for both human and mouse. The sample-specific kinase-substrate list, predicted using the Kinase Library, can also be integrated into the IncytrDB as validated interactions when applicable. **D.** The method overview. Incytr requires scRNA-seq (or snRNA-seq) data for one or more conditions, user-assigned cell cluster labels and user-selected gene lists as inputs. Additional data, such as proteomics, phosphorylation data, and predicted kinase-substrate relationships, can enhance analysis results and uncover novel signaling pathways. The core functions of Incytr include: (1) Inference of ligand-target (L-T) signaling pathways using IncytrDB and the input gene list; (2) Processing of proteomics and phosphorylation data for integration; (3) Identification of phosphorylation-related kinases using the Kinase Library predictions; (4) Quantitative analysis of inferred L-T pathways to assess pathway activity; (5) Evaluation of inferred L-T pathways using the Pathway Differential Score (PDS). **E.** Additionally, Incytr provides intuitive visualizations, including a circle plot for overall signaling between clusters, a UMAP plot for recognizing pathway patterns, and a river plot to highlight significant L-T pathways.

We extend this concept to multi-modal data by developing Incytr (Inference of Cell Signal Transmission), a method for efficient discovery of cell signaling pathways through integration of scRNA-seq, proteomics, phosphoproteomics, and kinase–substrate specificity data (Figure 1B). Incytr identifies cell-type-specific Ligand → Receptor → Effector Molecule → Target (L–R–EM–T) pathways from scRNA-seq, generalizable to

L-R-…-EM-…-T (for simplicity, we use L-R-EM-T throughout the text), as the PPI database includes indirect interactions mediated by associations or co-expression within the same pathway. It conditions gene co-expression on prior or inferred interaction knowledge, including sample-specific kinase–substrate relationships predicted by Kinase Library (KL) [Johnson et al., 2023; Yaron-Barir et al., 2024], and supports differential pathway analysis across conditions. The framework optionally incorporates additional modalities by estimating cell-type-specific proteomic and phosphoproteomic signals and integrating kinase activity predictions to refine pathway inference.

We demonstrate Incytr’s utility in COVID-19, Alzheimer’s disease, and cancer, where it recovers known pathways and generates novel, cell-type-specific hypotheses supported by multiple data types. Integration with biomarker and drug databases further enables target discovery. Finally, Incytr-derived pathways can serve as training data for simple NLP models, enhancing the analysis of cell–cell communication in health and disease.

## Results

### Overview of Incytr

Incytr efficiently identifies cell signaling pathways from scRNA-seq data alone or integrated with proteomics, phosphoproteomics, and kinase-substrate specificity (Figure 1, Supplementary Figure 1). The required inputs for Incytr are single-cell transcriptomics data, cell group labels, and user-selected sender and receiver genes which can be any genes measured in the data. The optional inputs include condition labels for the cells, proteomics and phosphoproteomics data, and a predicted kinase-substrate list (see Kinase-Substrate Matching in Supplementary Materials). With the input data, Incytr performs the tasks in the following modules:

*(1). Database construction and inference of the ligand–target signaling pathways.* The signaling pathway from the ligand to the target is considered to have the ‘L-R-EM-T’ signaling structure. Based on such a structure, the Incytr database (IncytrDB) is constructed, utilizing multiple data sources (Figure 1C, see Materials and Methods), and is available for human and mouse species. In addition, the predicted kinase-substrate list (optional input) can also be integrated into the IncytrDB as validated interactions when available. With the IncytrDB, we then infer all the ligands from the input sender genes, and the receptors, effector molecules, and targets from the given receiver genes, to ensure each pathway has the ‘L-R-EM-T’ signaling structure (Figure 1D, see Materials and Methods).
*(2). Quantitative analysis of the inferred pathways using the single-cell transcriptomics data.* For each condition, we first calculate the expression level of the genes in the inferred pathways, then calculate the statistical significance via permutation tests, and predict the signaling probability using a Hill function model (Figure 1D, see Materials and Methods). We then identify the differentially expressed pathways by calculating the adjusted fold change (aFC) value (see Materials and Methods).
*(3). Processing and the integration of multi-omics data.* We provide instructions for simulating cell-type-specific proteomic (or phosphoproteomics) data from scRNA-seq and bulk proteomics (or phosphoproteomics). Then, Incytr integrates the multi-omics data and quantifies the cross-condition change for each inferred pathway (Figure 1D, see Materials and Methods).
*(4). Identifying kinase-substrate relationships in the inferred pathways.* With the kinase-substrate predictions from Kinase Library [Johnson et al., 2023], Incytr first identifies signaling-involved kinases (SiKs) and signaling-related kinases (SrKs) from the ligand-target signaling pathways, then quantifies their relative phosphorylation activity in the corresponding cell group using the Exclusiveness Index (EI), based on the principle that a kinase substrate’s exclusively high expression in one cell group can indicate that such a group is undergoing a strong kinase-specific phosphorylation. Furthermore, Incytr calculates the SiK-score to evaluate how the phosphorylation activity supports the L-T pathway based on the SiKs’ EI (Figure 1D, see Materials and Methods).
*(5). Evaluating the inferred pathways based on multi-modal analysis.* A comprehensive quantitative evaluation of the inferred L-T pathways on their cross-condition difference based on the multi-modal analysis is performed by Incytr based on the following steps: (1) the transcriptomics-based pathway differential score (T-PDS) is calculated based on the aFC value on the signaling probabilities between conditions and used as the base score; (2) the proteomics and phosphorylation-based pathway differential score (P-PDS and Ph-PDS) based on the analysis using the proteomics and phosphorylation data, respectively; (3) the kinase-based pathway differential score (K-PDS) is calculated based on the aFC value of the SiK-score between conditions; (4) the pathway differential score (PDS) (the overall degree to which an L-T pathway is differential between conditions) is calculated by adding weighted P-PDS, Ph-PDS, and K-PDS to the base score (Figure 1D, see Materials and Methods).
*(6). Visualizing and exporting results via the Incytr web interface.* Incytr provides informative and interactive visualizations (Figure 1E) through the module Incytr-Viz (an open-source Python package). This module uses the Incytr analysis results as the input to view, filter, and explore the inferred signaling pathways on an interactive window. For example, the Incytr-Viz displays the number of pathways occurring between pairs of cell populations through a cellular interaction graph, and inspects the genes that comprise the pathways via a river (Sankey) plot (Figure 1E). The users can further filter the signaling pathways by applying different metrics, such as selecting specific genes or cell groups, assigning cutoff values to the PDS, etc. (see Materials and Methods).

### Incytr Validation: Biological Relevance and Benchmarking Against Existing Solutions

To validate that Incytr can identify meaningful signaling pathways based on causal relationships among ligand, receptor, effector, and target, we first applied Incytr to a published multimodal dataset (transcriptomics and proteomics) from a Th17 cell differentiation study [Ariss et al., 2024]. The study examined naïve CD4+ T cells from mouse spleens and lymph nodes, cultured under different stimulation conditions: Th0 (anti-CD3 + anti-CD28), non-pathogenic Th17 (npTh17; anti-CD3 + anti-CD28 + IL-6 + TGFβ), pathogenic Th17 (pTh17; anti-CD3 + anti-CD28 + IL-6 + IL-1β + IL-23), or PMA and Ionomycin (anti-CD3 + anti-CD28 + PMA + Ionomycin). Cells were collected at 0, 10, 45 minutes, 6 hours, and 24 hours post-stimulation.

We aimed to determine whether Incytr could recapitulate a key observation from the study regarding the role of Stat3 in Th17 differentiation. As previously shown, this process begins with IL-6 binding to its receptor, IL-6R, activating JAK kinases, which then phosphorylate Stat3. Phosphorylated Stat3 dimerizes and translocates to the nucleus, inducing genes essential for Th17 differentiation [Ariss et al., 2024]. The study identified Stat3’s direct gene targets, differentially regulated by pStat3 signaling in npTh17 and pTh17 conditions. As demonstrated (Supplementary Figure 2A,B), Incytr successfully identified these functional relationships between Stat3 and its target genes in both npTh17 and pTh17 conditions.

As another approach, we used a study with an oncology model, where A549 cells were treated with inhibitors targeting distinct cancer signaling nodes for 24 hours [McFarland et al., 2020]. Single-cell transcriptional response data from the study were analyzed using Incytr for differential pathway modulations based on aFC values (median aFC > 1 indicating pathways suppressed by drug treatment; refer to the methods section in Supplementary Materials).

The analysis recapitulated the expected mechanisms of action for several agents (Supplementary Figure 2C). The MCL1 inhibitor AZD5591 [Liu et al., 2021] and BET inhibitor JQ1 showed broad suppression across RTK-driven and NF-κB survival pathways, consistent with global transcriptional repression. Interestingly, JQ1 showed markedly lower UMI counts per cell, despite having the highest cell count, further supporting the transcriptional repression. Gemcitabine, a nucleoside analog, predominantly suppressed hypoxia and cell-cycle associated pathways, reflecting its impact on DNA synthesis and replication stress. The ATR/Chk1 inhibitor prexasertib strongly affected ubiquitin-mediated proteotoxic stress pathways, consistent with checkpoint disruption and replication stress. The MEK1/2 inhibitor trametinib suppressed KRAS-driven and MAPK cascade pathways, while the mTORC1 inhibitor everolimus primarily affected PI3K signaling and downstream survival pathways. In contrast, Incytr did not detect sufficient evidence of transcriptional suppression signatures for the RTK inhibitor afatinib or the PI3Kα inhibitor taselisib in this study to support a clear drug–pathway correlation. Overall, the Incytr-derived suppressed pathway signatures were well aligned with the known mechanisms of action of the tested drugs in this assay.

Incytr analysis of pathways upregulated following drug treatment (aFC < –1) revealed distinct drug-specific signaling adaptations (Supplementary Figure 2C). Gemcitabine upregulated RTK-associated signaling, while the ATR/Chk1 inhibitor prexasertib enriched Notch-like stemness programs. RTK inhibition (afatinib) and MEK inhibition (trametinib) were associated with increased immune-response pathways. In contrast, PI3K/mTOR inhibition (everolimus, taselisib), though weaker in signal, favored RTK/MAPK-mediated compensatory signaling. AZD5591 and JQ1 predominantly induced cell-cycle stress and RTK/hypoxia-stress signaling, respectively. Collectively, these pathway shifts may reflect adaptive or escape programs triggered by targeted drug pressure and warrant further validation.

The adaptive signaling responses identified in drug-treated A549 cells converge on pathways already recognized as therapeutic targets: EZH2-mediated chromatin remodeling (gemcitabine) [Li et al., 2021], RHO/cytoskeletal signaling (prexasertib) [Barcelo et al., 2023], CDK4/6-driven cell cycle compensation (afatinib) [Zhang et al., 2021], and interferon/immune pathway activation (trametinib) [Kang et al., 2019], each with corresponding agents in clinical development or under active investigation.

To further assess the biological relevance of Incytr-inferred signaling pathways, we leveraged the study of cytokine-mediated communications in immune cells [Cui et al., 2024], which profiled single-cell transcriptomes of over 17 immune cell types in response to 86 cytokines in vivo in mouse lymph nodes. Applying Incytr to the cytokine–cell type combinations reported in the original study (Method Validation Using the Mouse Immune Data in the Supplementary Materials), we found that approximately 90% of the Hallmark pathway IDs associated with Incytr-inferred cytokine-driven signaling pathways (i.e., where the selected cytokine serves as the ligand) were consistent with those reported in the original study (Figure 2A). Even when including the Hallmark pathway IDs mapped from all signaling pathways, not limited to those directly involving the cytokine molecule, about 60% of the Hallmark pathway IDs remained supported by the original study (Supplementary Figure 3). This supports the biological validity of Incytr’s inferred pathways and suggests they capture relevant ligand-receptor-mediated signaling responses.

**Figure 2.**
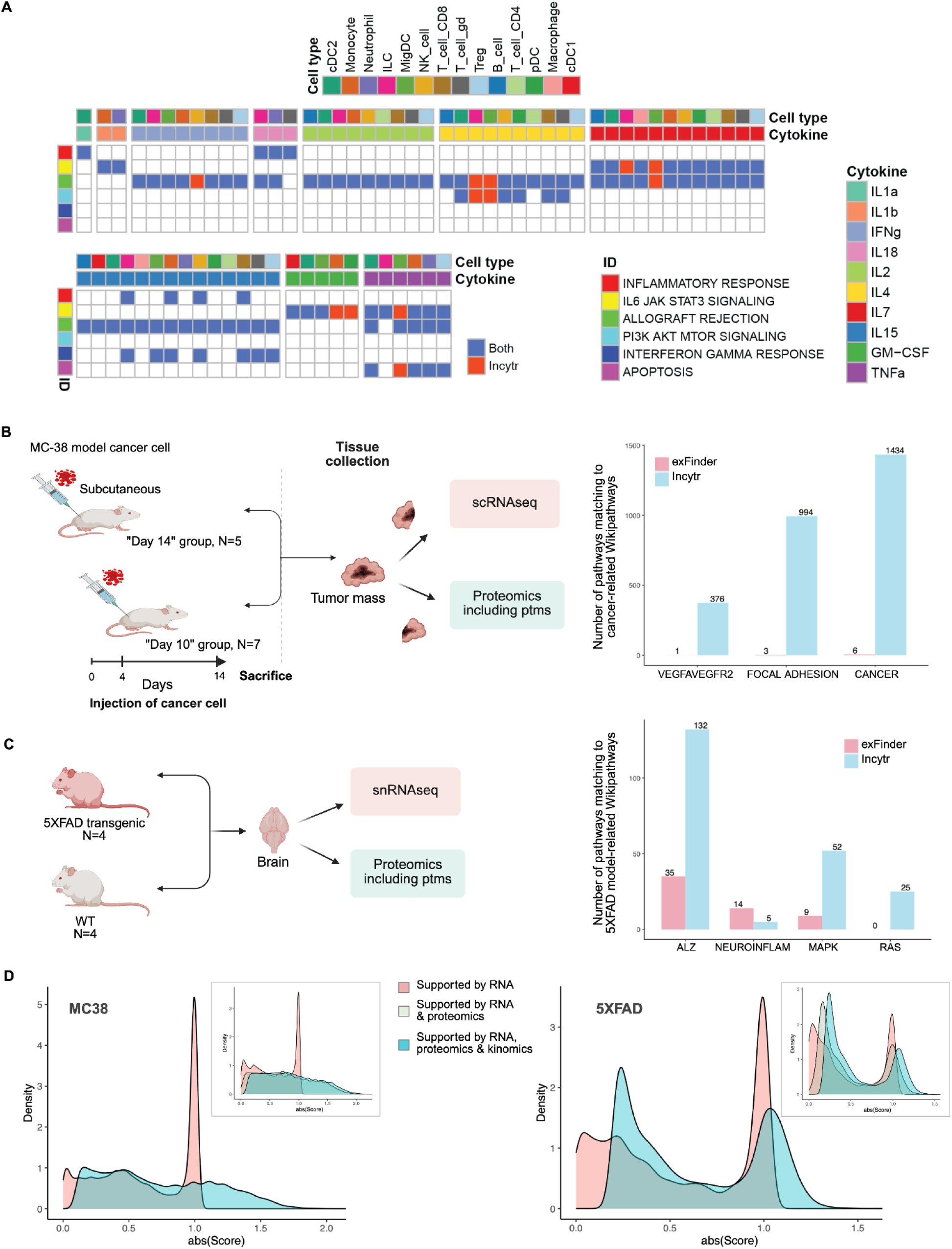
Incorporating multimodal data into the signaling pathway discovery process adds significant value. **A.** About 90% of the Hallmark IDs identified by Incytr in response to specific cytokine stimulations were also reported in the ‘cytokine dictionary’ study [Cui et al., 2024]. We present only the results for Cytokine → R → EM → T networks, where the cytokine matches the one used for stimulation. In the plot, blue indicates Hallmark IDs identified by both Incytr and the cytokine dictionary study, while red indicates those identified by Incytr only. Cartoon representation of the “MC38” (**B**) and “5XFAD” (**C**) study designs, along with the distribution of exFINDER- and Incytr-discovered signaling pathways mapped to cancer-specific and 5XFAD model-specific WikiPathways. Paths were counted for Incytr if they had a p value < 0.05 in at least one condition and a minimum PDS of 0.2, and paths were counted for exFINDER if they had a minimum score (calculated using the same scoring function is Incytr) between conditions of 0.2. Paths were aggregated across all pairwise sender-receiver group combinations. **D.** The added value of incorporating proteomics and kinomics data in pathway scoring lies in its ability to enhance confidence in pathway identification. The inclusion of multimodal data not only highlights the “differential” pathways (with a score shift above 0.76, corresponding to a one-fold change difference between groups/conditions) but also “rescues” pathways with borderline significance (shifting scores from 0 toward positive values). The inset illustrates the decoupled contributions for RNA alone, RNA & proteomics, and RNA & proteomics & kinomics. Only pathways that include all three modalities (approximately 35% in the MC38 dataset and 55% in the 5XFAD dataset) are shown in the inset.

We next assessed whether Incytr can recapitulate known signaling pathways relevant to the biological systems analyzed in this study, including mouse models of colon cancer and Alzheimer’s disease, as well as patient samples from COVID-19 cohorts. To this end, we inferred four-step signaling cascades (ligand-receptor-effector-target) from RNA-seq and multi-modal data and mapped them onto the WikiPathways database (https://www.wikipathways.org/). This analysis showed that Incytr successfully recapitulates pathways specific to the MC38 and 5XFAD mouse models, as well as COVID-19–associated signaling (Figure 2B, C; Supplementary Data 1).

To benchmark this performance, we compared Incytr with GSEA and exFINDER by mapping statistically significant pathways identified by each method to WikiPathways across the same datasets. This comparative analysis (Figure 2B, C, Supplementary Data 1) demonstrates that Incytr more effectively rediscovers known cancer-, 5XFAD-, and COVID-specific signaling pathways.

Integrating additional modalities, such as proteomics, phosphoproteomics, and kinomics, directly demonstrates the added value of multimodal analysis for pathway identification. By leveraging complementary data types, Incytr prioritizes “top-tier” signaling networks supported across modalities and recovers biologically relevant pathways that are often missed or weakly supported by RNA-seq alone (Figure 2D, Supplementary Figure 4). Consistent with this, we observe a substantial increase in the number of identified pathways when proteomics and kinomics data are incorporated into the input gene lists, as well as when the Incytr protein-protein interaction (PPI) network is expanded using dataset-specific substrate-kinase (KL) predictions (Supplementary Figure 4).

To further illustrate the utility of the multimodal approach, we applied Incytr to paired snRNA-seq and snATAC-seq data from PBMCs of individuals with a chronic viral infection (people with HIV virally suppressed with antiretroviral therapy), focusing on L-R-EM-T signaling in CD8⁺ T cells, given their importance in antiviral immune response. Incytr analysis of the snATAC-seq data revealed multiple signaling networks within a prominent NK→CD8⁺ T cell interaction axis. Specifically, NK cell-derived MHC-II components (HLA-DMA/DMB/DPB1) were predicted to engage CD4 on CD8 T cells dually expressing CD4, inducing antiviral programs characterized by upregulation of MX1/2, IFIT1/3, and IFITM1 (Supplementary Figure 4D). Notably, these signals were not detected in scRNA-seq data alone, likely due to transient or low expression levels (Supplementary Figure 4D).

To validate these findings, we turned to orthogonal CyTOF measurements, where we implemented a recently developed and validated viral sensing/restriction panel [George et al., 2025] on the same specimens, selecting key antiviral factors (MX1, MX2, AIM2, IFIT3) identified from our paired snRNA-seq/snATAC-seq Incytr analysis. We confirmed elevated protein levels of these antiviral effectors in CD4⁺CD8⁺ double-positive T cells compared to CD8⁺ T cells that did not co-express CD4 (Supplementary Figure 4E). Of note, NK cell expression of MHC-II machinery [Ustiuzhanina et al., 2023; Luo et al., 2017] and the involvement of CD4⁺CD8⁺ T cells [Zahran et al., 2021; Riou et al., 2021; Zhang et al., 2022] in antiviral responses have previously been reported in viral infections, including not only by HIV, but by Hantaan virus and SARS-CoV-2. Together, these results demonstrate that Incytr’s multimodal integration framework, particularly through the incorporation of chromatin accessibility data, can uncover biologically meaningful signaling interactions that are not apparent from transcriptomic data alone.

### Incytr Suggests Hypotheses for How Fibroblasts Can Influence Cancer Cells to Undergo Epithelial-Mesenchymal Transition (EMT)

A high-level comparative analysis between the Day 10 and Day 14 MC38 mouse model sample groups revealed notable differences in the frequency of EMT and cancer cells (Figure 3A). This suggests that a significant portion of the cancer cell population acquires an EMT phenotype in preparation for metastasis on Day 14 as compared to Day 10. We then asked which cell-cell communication patterns might be relevant to this transition.

**Figure 3.**
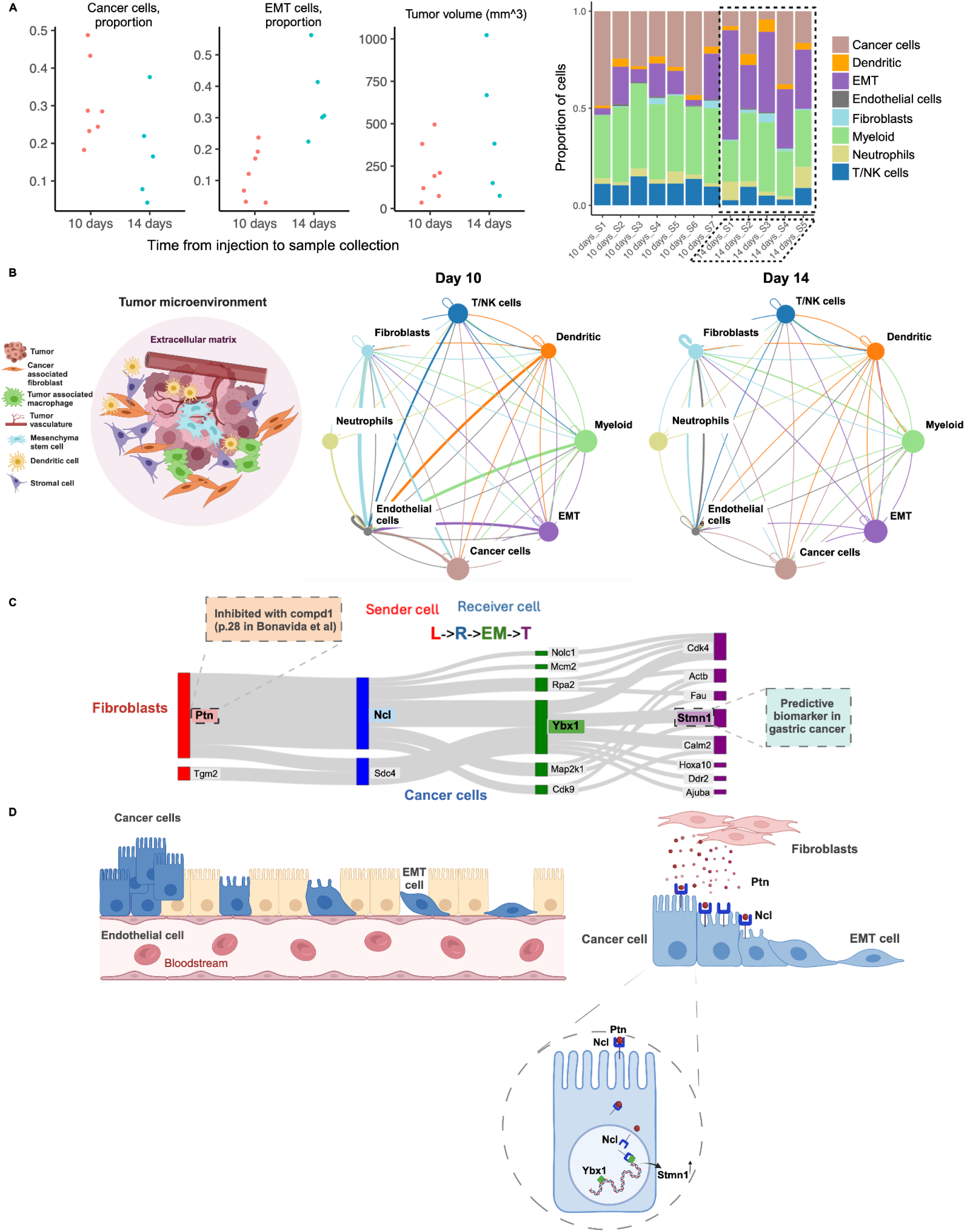
Incytr offers a holistic view of how cell-cell communication patterns change as cancer progresses, enabling the generation of novel hypotheses that extend beyond known cell signaling pathways. **A.** Per-sample cell composition analysis revealed a significant increase in the EMT cell compartment by day 14. **B.** The schematic representation of the tumor microenvironment (TME) illustrates its complex and heterogeneous cell composition, highlighting the need to consider cross-talk between multiple cell types simultaneously, rather than adopting a one-cell-type-centric approach. Statistically significant pathways with more than a two-fold change difference between the groups (signaling probability > 0.25, T-PDS < -0.76, and PDS < -0.76) are shown for all possible pairwise cell-cell interactions in the Day 10 and Day 14 groups. The area of each circle is proportional to the log of the size of one cell population, while the thickness of the connected edges indicates the number of discovered highly scored pathways. **C.** A subset of Day 10 signaling pathways discovered by Incytr (p-value<0.05, signaling probability>0.25, T-PDS<-0.96, and PDS<-0.76) involve fibroblasts sending signals to cancer cells via the Ptn-Ncl ligand-receptor interaction. The following annotations were extracted from the Liceptor and Biomarker databases: 189 small molecule compounds identified for targeting Ptn. For example, compound 1 (page 28) from [Bonavida (Nereus Pharmaceuticals, Inc.), US Patent Application 2012/0282168 A1], with the following details: function: inhibitor; therapeutic data: ovarian cancer, breast cancer, colon cancer, lung cancer, multiple myeloma; molecular structure (SMILES): CC12OC(=O)C1(C(O)C1C=CCCC1)NC(=O)C2CCCl. Similar annotations are available for Ncl and Ybx1 in the Liceptor database (not shown here). Stmn1 identified as a predictive biomarker- clinical significance: patients with Stmn1 overexpression showed poor survival with docetaxel treatment; molecular alteration: overexpression; investigation technique: immunohistochemistry. **D.** Hypothesized interaction: Ptn binds to Ncl on the cancer cell surface, and the complex is internalized. Ncl then transports Ptn to the nucleus and cytoplasm. In the nucleus, Ncl interacts with Ybx1, modulating its activity. Ybx1, in turn, regulates the expression of Stmn1 and other genes, driving changes associated with EMT, cell proliferation, and migration. The upregulation of Stmn1 and other Ybx1 target genes enhances the cancer cell’s ability to proliferate, migrate, and invade, thereby contributing to cancer progression and metastasis.

To explore this, we characterized each sample by the presence of eight cell types, including cancer cells, EMT cells, endothelial cells, fibroblasts, and key immune cell types (Figure 3A). We applied Incytr to analyze highly expressed and differentially expressed genes (from scRNA-seq and proteomics data) across the conditions and cell types, comparing cell signaling pathways between the Day 10 and Day 14 groups. Several unique signaling pathways emerged for each group (Figure 3B). We focused on the most prominent differential axis of interaction with cancer cells on Day 10, specifically the interaction with fibroblasts.

For illustrative purposes, we highlight one of the statistically significant pathways with more than a two-fold change difference between the Day 10 and Day 14 groups: the interaction between fibroblasts and cancer cells via the Ptn-Ncl ligand-receptor pair. The interaction between pleiotrophin (Ptn), nucleolin (Ncl), Y-box binding protein 1 (Ybx1), and stathmin 1 (Stmn1) in cancer cells forms a complex signaling pathway (Figure 3C, D, Supplementary Figure 5A, Supplementary Data 2) that influences critical cellular processes, including proliferation, migration, and epithelial-mesenchymal transition (EMT). Ptn, a heparin-binding growth factor secreted by fibroblasts, binds to Ncl on the surface of cancer cells [Koutsioumpa et al., 2013; Lamprou et al., 2022]. This interaction facilitates the internalization of the Ptn-Ncl complex [Wang, 2020]. Once internalized, Ncl transports Ptn to different cellular compartments, including the nucleus and cytoplasm, triggering various signaling pathways. In the nucleus, Ncl is involved in ribosome biogenesis and gene expression regulation, while in the cytoplasm, it plays a role in mRNA stabilization and translation [Abdelmohsen & Gorospe, 2012].

Ybx1 acts as a transcription factor regulating genes involved in cell proliferation, apoptosis, and EMT. Ncl can modulate Ybx1 activity [Ke et al., 2021], enhancing its role in promoting oncogenic processes, including the expression of Stmn1 (Supplementary Data 2). Stmn1, a microtubule-destabilizing protein, is crucial for cell cycle progression and migration. Ybx1-mediated upregulation of Stmn1 increases cancer cell motility and invasiveness, contributing to metastasis [Cai et al., 2022].

A potential therapeutic strategy would target key points in the Ptn-Ncl-Ybx1-Stmn1 (Supplementary Data 2) signaling pathway. Disrupting the ligand-receptor interaction (Ptn-Ncl), transcriptional regulation (Ybx1), or cytoskeletal dynamics (Stmn1) presents several avenues to reduce cancer cell motility, invasiveness, and metastasis. Combination therapies or nanotechnology-based approaches could enhance the efficacy of these treatments, particularly in tumors reliant on this pathway. As shown in Figure 3C, overlaying data from the Liceptor and Biomarker databases provides immediate information about small molecule compounds previously used to target individual components in the Ptn-Ncl-Ybx1-Stmn1 signaling pathway, indicating whether any of these molecules have been reported as biomarkers, thereby informing potential treatment strategies.

### A Holistic View of Cell Signaling in the 5XFAD Alzheimer’s Disease Mouse Model Suggests New Hypotheses About Neuroprotective Pathways

The 5XFAD model [Wang et al., 2015; Oakley et al., 2006] is a widely used transgenic mouse model for Alzheimer’s disease (AD), carrying five familial AD mutations—three in the amyloid precursor protein (APP) and two in the presenilin 1 (PSEN1) genes. These mutations lead to aggressive amyloid-beta (Aβ) plaque formation, mimicking key features of human AD.

To better understand the inflammatory and immune-related signaling in AD, we applied Incytr to multi-modal data (mRNA-seq, proteomics, phosphoproteomics, kinomics) from 5XFAD mouse model samples (Figure 4). Since this model is often used to study immune responses, particularly microglia-regulated neurodegeneration [Wang et al., 2015; Zhou et al., 2020], we focused on signaling pathways involving microglial cells. First, we verified that Incytr successfully recapitulates known signaling cascades involving microglia in the 5XFAD mouse model (Figure 4A,B), which may influence neuroinflammation, Aβ clearance, and overall microglial function.

**Figure 4.**
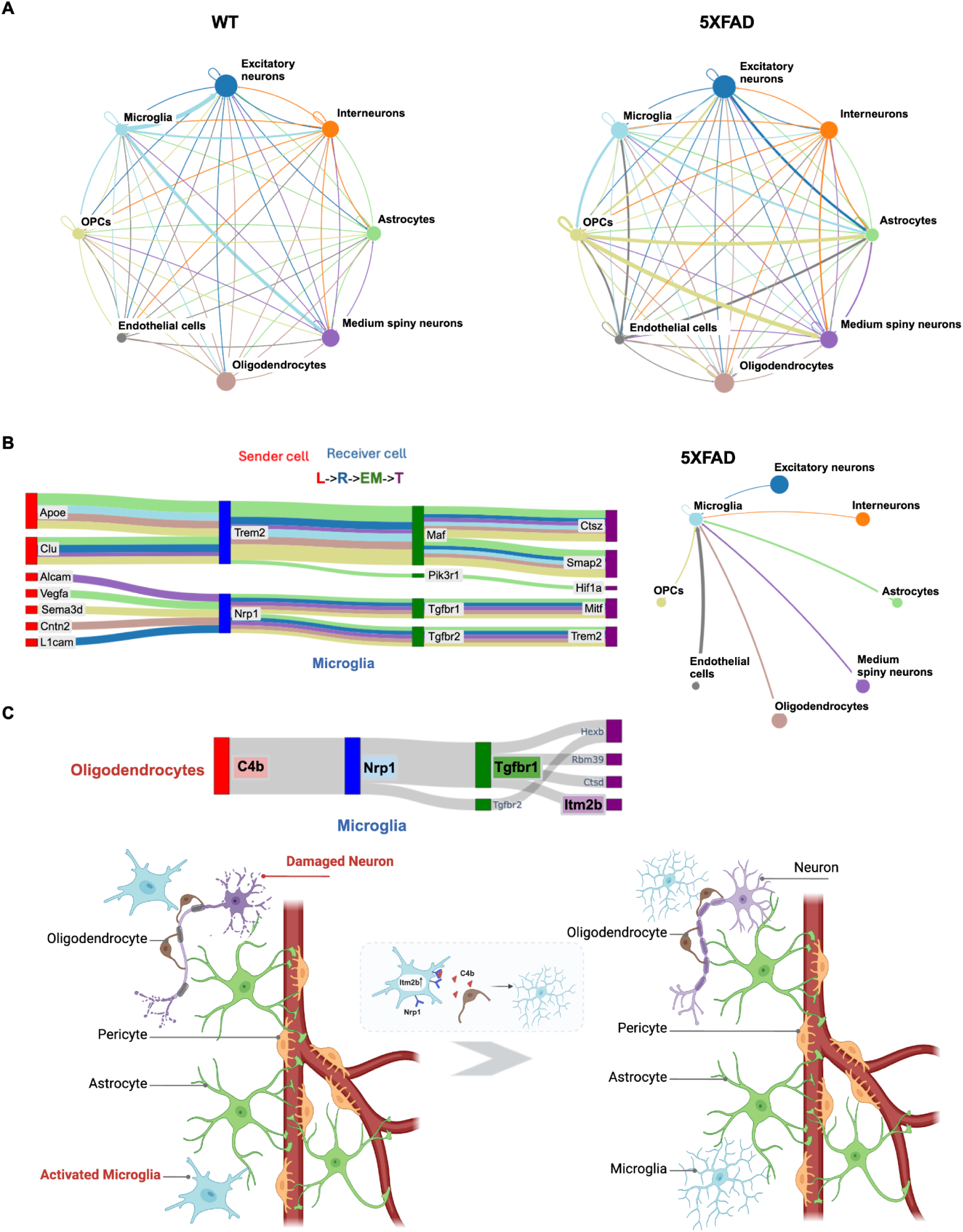
As suggested by Incytr, one potential pathway to achieve an anti-inflammatory state in microglia, leading to neuroprotection in the 5XFAD model, is through the C4b-Nrp1 ligand-receptor interaction between oligodendrocytes and microglia. **A.** Cell signaling patterns (signaling probability>0.25) with a fold change greater than 2 between wild type and 5XFAD mouse groups. **B.** Incytr has rediscovered signaling pathways that align with well-established mechanisms of cell-cell interactions in the context of neuroinflammation and neurodegeneration in the 5XFAD model. The Sankey plot is color-coded by sender group type. **C.** C4b released from oligodendrocytes can bind to the Nrp1 receptor on microglia. Upon C4b binding to Nrp1, downstream signaling through Tgfbr1 may occur. Activation of Tgfbr1 can lead to the expression of various target genes, including Itm2b. This signaling pathway can modulate microglial inflammatory responses, either promoting anti-inflammatory effects or exacerbating inflammation, depending on the context. The illustration depicts a neuroprotective scenario.

Among these pathways are the communications between oligodendrocyte precursor cells (OPCs), medium spiny neurons, and excitatory neurons with microglia via the Apoe (apolipoprotein E)-App (amyloid precursor protein), Apoe-Trem2 (triggering receptor expressed on myeloid cells 2), and Apoe-Lrp1 (low-density lipoprotein receptor-related protein 1) pathways (Supplementary Data 3). These pathways highlight how neuronal and glial interactions modulate microglial activity through Apoe, App, Trem2, and Lrp1 [Krasemann et al., 2017; Lin & Holtzman 2024; Shinohara et al., 2017].

The Apoe-App interaction is crucial for the metabolism and clearance of Aβ, a key pathological hallmark of AD. Apoe influences Aβ aggregation and deposition by affecting App processing and Aβ clearance [Lin & Holtzman 2024]. The Apoe-Trem2 axis plays a significant role in modulating microglial responses to Aβ and other neuroinflammatory signals. Trem2, expressed on microglia, interacts with Apoe to regulate phagocytosis, survival, and inflammatory responses of microglia [Yeh et al., 2016; Lin & Holtzman 2024]. Additionally, the Apoe-Lrp1 pathway is essential for lipid transport and Aβ clearance across the blood-brain barrier. Lrp1 binds to Apoe-containing lipoproteins and facilitates the endocytosis and degradation of Aβ [Tachibana et al., 2019; Shinohara et al., 2017]. These findings underscore the complex cellular crosstalk in the AD brain and demonstrate Incytr’s capability to elucidate critical signaling pathways that are well-established in AD pathology. By successfully identifying these known pathways, Incytr validates its effectiveness in capturing key molecular interactions that could be targeted for therapeutic intervention.

Building upon these results, we explored novel, high-scoring microglial pathways proposed by Incytr. Notably, we identified a previously uncharacterized pathway involving oligodendrocyte-released complement component 4b (C4b), which binds to Neuropilin 1 (Nrp1) receptors on microglia (Figure 4C). This interaction potentially triggers downstream signaling through transforming growth factor beta receptor 1 (Tgfbr1), initiating a cascade that modulates microglial activity. In microglia, Nrp1 is known to promote M2 polarization, enhancing phagocytosis of cellular debris and contributing to an anti-inflammatory environment. It is also involved in interactions with regulatory T cells, triggering the release of transforming growth factor beta (TGF-β), which leads to immunosuppression [Chuckran et al., 2020]. The Nrp1-mediated enhancement of microglial phagocytosis may facilitate the removal of Aβ plaques in AD, and impaired Nrp1 function could lead to decreased clearance and increased plaque burden.

This novel C4b-Nrp1 signaling pathway represents a potential mechanism by which oligodendrocytes influence microglial function and neuroinflammation in AD. Activation of this pathway can lead to the expression of genes such as Itm2b, which modulates microglial inflammatory responses. Depending on the context, Itm2b can promote anti-inflammatory pathways or exacerbate inflammation (Figure 4C, Supplementary Figure 5B). Mutations in the Itm2b gene are associated with familial British and Danish dementias, highlighting its role in neurodegenerative diseases [Vidal et al., 1999]. Moreover, Itm2b (also known as Bri2) has been shown to interact with Trem2, influencing Trem2 processing and expression levels [Del-Aguila et al., 2019; Yin et al., 2024]. In the 5XFAD mouse model, increased expression of Itm2b in microglia may play crucial roles in modulating inflammation, clearing Aβ, and enhancing neuroprotection, thereby contributing to the overall function of microglia in AD. All other potential direct influencers on the Itm2b gene expression in microglia could also be explored by Incytr, as shown on Supplementary Figure 6.

The identification of the C4b-Nrp1-Tgfbr1-Itm2b pathway (Supplementary Data 2) suggests new therapeutic opportunities. Since the interaction between C4b and Nrp1 initiates a signaling cascade that regulates microglial activity, therapeutic agents such as small molecules or monoclonal antibodies could be developed to modulate this interaction. By inhibiting or enhancing specific components of the pathway, it may be possible to shift microglial activation toward a more neuroprotective, anti-inflammatory state, potentially reducing harmful neuroinflammation in AD.

### Incytr Reveals the Differential Effects of Dexamethasone Treatment on T Cells Based on Their Phenotype in COVID-19 Patients

COVID-19 patients with severe symptoms or those requiring oxygen support are frequently treated with Dexamethasone (DEXA), a corticosteroid [The RECOVERY Collaborative Group, 2021]. DEXA’s potent anti-inflammatory properties help suppress the hyperactive immune response, or “cytokine storm,” often observed in severe COVID-19 cases. A central aspect of DEXA’s immune modulation is its impact on various T cell subsets, including regulatory T cells (Tregs), effector T cells (CD4+ and CD8+), and exhausted T cells. By reducing hyperactivation in effector T cells, DEXA minimizes tissue damage [Giles et al., 2018; O’Garra & Barrat, 2003].

DEXA’s effects are mediated by the glucocorticoid receptor, NR3C1. Signaling through NR3C1 varies between T cell phenotypes: for example, in effector T cells, DEXA inhibits pro-inflammatory cytokine production and proliferation, potentially reducing harmful inflammation [O’Garra & Barrat, 2003]. In Tregs, DEXA may enhance immunosuppressive functions via NR3C1, promoting immune balance. For exhausted T cells, DEXA may modulate exhaustion pathways, particularly in cells expressing high levels of PD-1 and TIM-3. The differential NR3C1 signaling across T cell phenotypes in COVID-19 patients treated with DEXA has practical implications for clinical management and drug development.

A deeper understanding of DEXA’s modulation of T cell signaling through NR3C1 could help optimize dosages in patients with varied immune dysregulation levels. Insights into this pathway could also inform combination therapies, pairing DEXA with immunomodulators or antivirals to reduce inflammation while preserving immune responses. Understanding the differential effects of DEXA on Tregs versus effector T cells could guide personalized treatment approaches tailored to specific immune profiles.

This understanding motivated us to apply Incytr to infer NR3C1 signaling patterns across T cell subtypes in COVID-19 patients treated with DEXA (Figure 5A, B). Our analysis shows that while most signaling patterns are not differential between Tregs, T effector memory, and naive T cells, some phenotype-specific signaling pathways are present (Figure 5C-E, Supplementary Figure 5C, Supplementary Figure 7, Supplementary Data 2). For example, DEXA-NR3C1-FOXP3-VIM, DEXA-NR3C1-FOXP3-NFKBIA and DEXA-NR3C1-FOXP3-BTG2 pathways are distinct in Tregs, reflecting their specialized role in immune suppression under inflammatory conditions like COVID-19.

**Figure 5.**
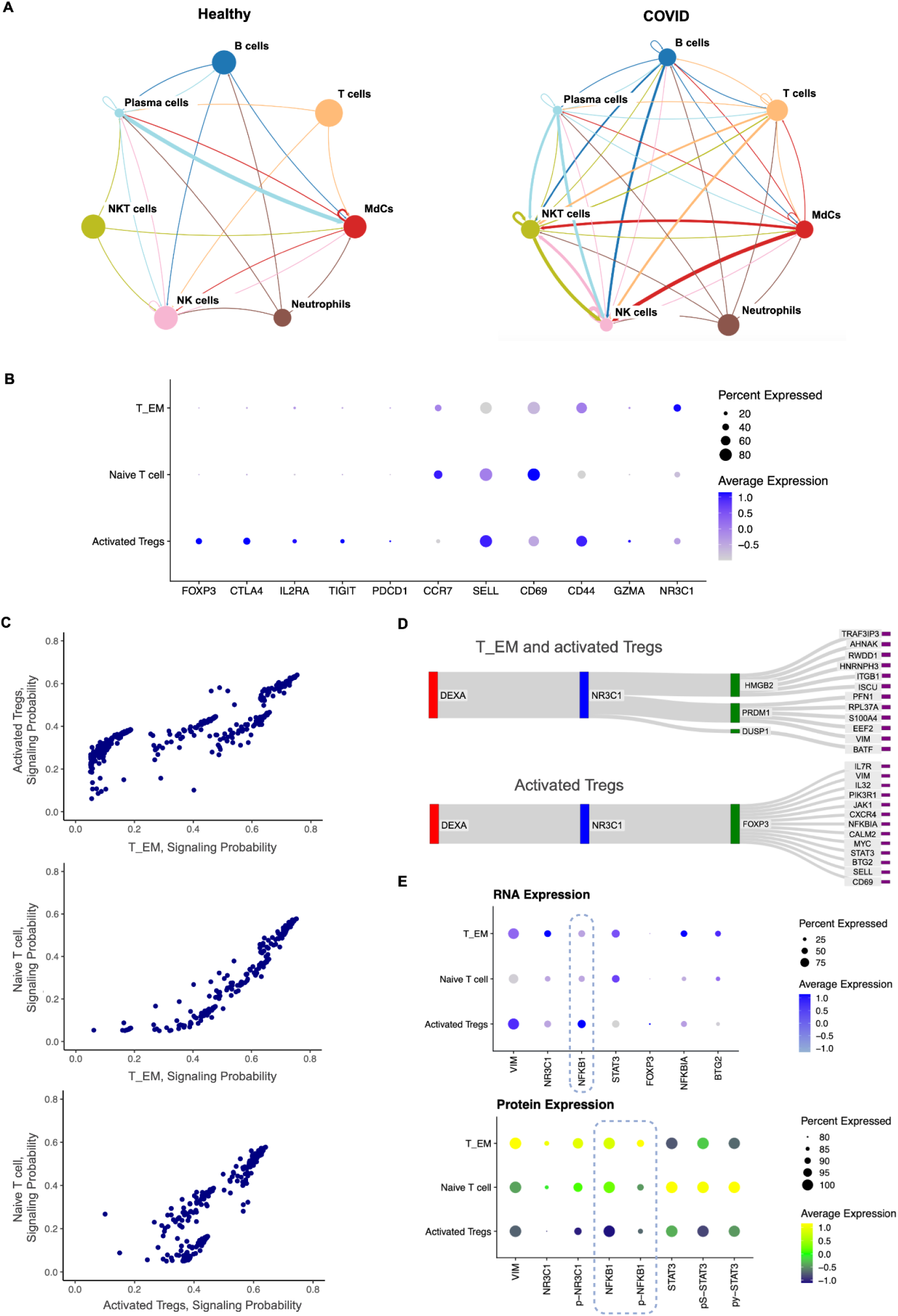
Incytr identifies T cell phenotype-specific signaling pathways in COVID-19 patients in response to Dexamethasone treatment. **A.** Cell signaling patterns (signaling probability>0.25) with a fold change greater than 2 between healthy controls and COVID-19 patient groups. **B.** Further subclustering of the T cell population in the primary COVID-19 patient cohort reveals three phenotypically distinct groups: regulatory T cells (Tregs), naive T cells (including both CD4+ and CD8+ T cells), and effector memory T cells (T_EM). **C.** Most of the Incytr-inferred signaling pathways relevant to DEXA treatment (with DEXA as the ligand) show no significant differential signaling pathways between Tregs, naive T cells, and T_EM. In the plot, each dot represents one L-R-EM-T signaling pathway. Additional rows were added to the transcriptomic expression matrix to simulate an artificial cell type X of one cell per condition with nonzero expression of DEXA (all other cells had 0 expression of DEXA, and X had 0 expression of any other genes). Incytr was run with X as the sender group. **D.** Representative groups of signaling pathways uniquely present in Tregs, as well as those shared between Tregs and T_EM cells, are highlighted. Analysis is shown for the first cohort of COVID-19 patients. An independent validation analysis of these discovered pathways was done using samples from the second cohort of COVID-19 patients. **E.** InTraSeq results for the second cohort of COVID-19 patients (N=3). The top panel shows mRNA expression, and the bottom panel displays protein expression.

When NR3C1 activates FOXP3 in Tregs, it reinforces their regulatory capacity—an effect not seen in non-Treg cells, where FOXP3 is either absent or does not play a similar role. VIM (Vimentin), involved in cytoskeletal dynamics, may support Tregs’ mobility and infiltration into inflamed COVID-19 tissues, aiding targeted suppression of inflammation. This contrasts with other cells, where Vimentin mainly supports structural stability rather than immune modulation. Knowledge of how VIM supports Treg mobility suggests a potential to enhance Treg infiltration in inflamed tissues through targeted modulation of the cytoskeletal pathways. This could be particularly relevant in preventing organ damage in severe COVID-19 cases, by enabling Tregs to suppress local inflammation more effectively.

NF-κB is a crucial transcription factor in inflammatory responses, playing a complex role in immune regulation. Its function in Tregs is particularly nuanced, especially in the context of COVID-19 and other inflammatory conditions. The suppression of NF-κB in Tregs appears to be a key mechanism for maintaining their anti-inflammatory role in inflamed environments, such as those seen in COVID-19. This suppression helps Tregs control cytokine storms without significantly compromising other immune functions. There are several mechanisms to regulate the NF-κB either at the level of transcription, translation or via NF-κB protein inhibition or degradation. The data presented (Figure 5E, Supplementary Figure 5C, Supplementary Figure 7) suggests that in Tregs, unlike T_EM and Naive T cells, NF-κB either doesn’t translate into protein or gets degraded.

While inhibiting NF-κB in Tregs might seem beneficial for controlling inflammation, it is important to consider the potential adverse effects in the context of COVID-19 or other inflammatory conditions. Although NF-κB is classically linked to pro-inflammatory responses, it is essential for Treg development, survival, and stability. In Tregs, NF-κB maintains their regulatory function, allowing effective suppression of excessive immune responses. Possible adverse effects of NF-κB inhibition in Tregs include increased risk of autoimmunity and impaired tissue repair, potentially slowing recovery in COVID-19 patients or leading to long-term damage in affected tissues, such as the lungs. Selective NF-κB inhibition could potentially prevent tissue damage in COVID-19 while minimizing risks like autoimmunity or compromised tissue repair.

### Semantic Analysis of Signaling Pathways Inferred by Incytr

From a signal transmission or communication perspective, the four-step (L-R-EM-T) pathways discovered by Incytr can be viewed as short sentences or phrases that cells use to communicate with each other. Therefore, the latest advancements in natural language processing are conceptually relevant for gaining a deeper understanding of this “cellular language”.

To this end, we applied the Doc2Vec model (https://radimrehurek.com/gensim/models/doc2vec.html), training it independently on each dataset (MC38, 5XFAD, COVID), where each gene name in a four-step pathway was treated as a word and the entire pathway as a sentence. We then extracted the embeddings and concatenated them with quantitative information (Figure 6A), such as the fold-change differences in signaling probabilities between conditions (e.g., 5XFAD vs WT).

**Figure 6.**
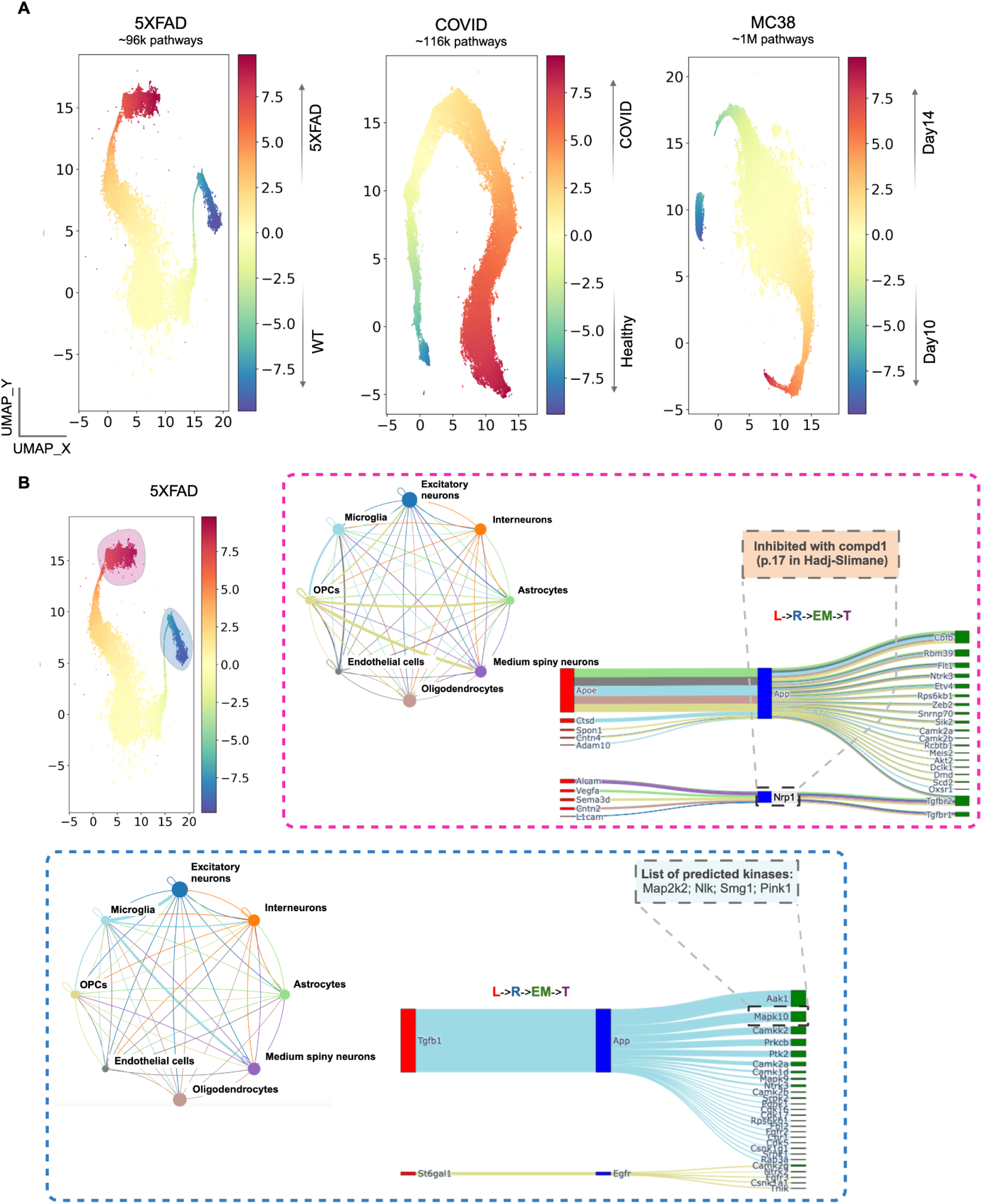
Semantic embeddings concatenated with quantitative fold-change data enable the identification of condition-specific signaling patterns. **A.** Embeddings for each dataset were color-coded based on the adjusted log2 fold-change between conditions, calculated using signaling probabilities for each L-R-EM-T pathway. Colors indicate signaling probability differences between conditions. **B.** An example of emerging pattern characterization: the red subset represents pathways that share similar semantics and are most probable in 5XFAD mice, while the blue subset shows those most probable in WT mice. In the red inset: an illustration of molecule annotations extracted from the Liceptor database, showing a small molecule compound identified for targeting the Nrp1 receptor. For example, compound 1 (page 17) from [Hadj-Slimane (Tragex Pharma), EP 2823816 A1, 2015], with the following details- function: inhibitor; therapeutic data: bladder cancer, thyroid cancer, liver cancer, atherosclerosis, rheumatoid arthritis; organism: Human (MDA-MB-231 cells); molecular structure (SMILES): CCOc1ccccc1NC(=O)c1ccc(C)c(S(=O)(=O)Nc2ccc(C)cc2)c1. In the blue inset: an illustration of kinases predicted by the Kinase Library to phosphorylate the binding sites on Mapk10. The Sankey plots are color-coded by sender group type.

This analysis allowed us to extract both semantic and numerical insights from signaling pathways. The Doc2Vec model captures the relationships between genes (words) based on their co-occurrence within the signaling pathways (sentences). This allows for uncovering patterns in gene-gene interactions that may be invisible through traditional analysis, potentially identifying functionally related genes that work together in pathways. These semantic embeddings can help identify novel gene functions or classify genes into groups based on shared signaling roles. When we concatenate the embeddings with quantitative fold-change data, the model can learn how changes in gene expression levels (e.g., 5XFAD vs WT) are associated with particular signaling pathways. This helps in identifying condition-specific signaling dynamics—for instance, how specific signaling pathways behave differently in disease versus healthy states (Figure 6B).

The embeddings capture the “meaning” of signaling pathways, while the quantitative data adds a contextual layer (e.g., how active or suppressed a pathway is in certain conditions). This combination allows us to infer which groups of genes or pathways are more prominent under certain biological conditions. It can also help predict how modifications in one part of the pathway (e.g., ligand-receptor interaction) might affect downstream targets and the overall biological function. Comparing the embeddings across different datasets (like COVID vs cancer models), could help determine shared signaling pathways across diseases or find specific signaling modules that may serve as biomarkers for particular conditions. Integration with databases of previously characterized drug molecules and biomarkers can provide insights into potential treatment options across various disease settings (Figure 6B, Supplementary Figure 8).

## Discussion

The ability to systematically uncover cell signaling pathways from multi-modal data offers transformative benefits. Integrating diverse data types (e.g., transcriptomics, proteomics, phosphoproteomics, kinomics) provides a more comprehensive view of cellular function and regulatory mechanisms across different biological layers. As shown, multi-modal data enables cross-validation across data types, resulting in more accurate and robust predictions of signaling pathways. This approach provides corroborative evidence for the same event-a signaling pathway-enhancing confidence in the findings. For example, mRNA abundance from transcriptomics alone is insufficient to infer protein abundance [Chen et al., 2020; Reimegård et al., 2021], and proteomics does not fully capture protein modifications that significantly impact protein structure and function [Zhong et al., 2023; Naowarojna et al., 2021]. Since diseases often involve disruptions across multiple pathways at different stages of signal transmission (e.g., transcription, translation), multi-modal data provides a more holistic understanding of these disruptions, enabling accurate modeling of disease mechanisms and progression.

Incytr offers a valuable framework for discovering cell signaling pathways by integrating scRNA-seq, proteomics, phosphoproteomics, and kinomics data. The method is designed to incorporate additional data modalities as they become available. An important enhancement could be integrating spatial information to refine pathway inference, particularly for cell-cell interactions requiring immediate sender-receiver proximity. Tools such as CellPhoneDB (V3) [Garcia-Alonso et al., 2021], COMMOT [Cang et al., 2023], and NeST [Walker & Nie, 2023] can support this refinement.

Our application of Incytr to COVID-19 patient data, MC38, and 5XFAD mouse model data has recapitulated known biological system-specific signaling pathways such as cancer specific pathways, Alzheimer’s disease specific pathways such as Apoe-App, Apoe-Trem2, and Apoe-Lrp1. Additionally, it uncovered novel interactions, including the C4b-Nrp1 pathway and other previously unknown pathways, that may play significant roles in disease pathology. These findings highlight Incytr’s capacity to identify critical signaling pathways and generate new hypotheses about pathways that could be targeted for therapeutic intervention.

It is important to note that signaling pathways inferred by Incytr, exFINDER, NicheNet, and similar tools are primarily based on associations rather than direct causal links. These methods use correlation-based approaches to suggest potential ligand–receptor interactions and downstream effects from gene expression data. While they do not inherently establish causality, we demonstrate that Incytr’s inferred pathways are enriched for biologically meaningful, causally linked interactions. This is partly due to Incytr’s design, which restricts candidate networks to L–R, R–EM, and EM–T pairs supported by empirical evidence, either direct interactions or co-membership in known pathways, thus increasing the likelihood of true causal relationships. Nonetheless, experimental validation or orthogonal methods, such as perturbation assays, remain essential to confirm causality.

As shown here, Incytr provides a systematic approach to inferring signaling pathways and enables the generation of large-scale, meaningful training data. This approach enhances our ability to understand fundamental biological mechanisms and supports practical applications, such as target and drug discovery, and beyond.

## Materials and Methods

### Incytr Overview

The Incytr computational method (Figure 1, Supplementary Figure 1, Supplementary Table 1) performs six key functions: (1) Inferring ligand-target signaling pathways from input gene lists using prior knowledge of protein interactions from the Incytr database (IncytrDB) (Figure 1C). (2) Quantitatively analyzing the inferred signaling pathways under different conditions using single-cell transcriptomics data (Figure 1D). (3) Processing and integrating multi-omics data to enhance pathway analysis (Figure 1D). (4) Identifying kinase-substrate relationships in the inferred pathways from the Kinase Library. (5) Evaluating the inferred pathways based on multi-modal analysis (Figure 1D). (6) Visualizing and exporting results via the Incytr web interface (Figure 1E).

### Incytr Database Integration

A database that captures the “ligand-receptor-effector molecule-target” structure is essential for inferring signaling pathways (Figure 1C). We built IncytrDB by integrating multiple publicly available datasets, including the exFINDER database (exFINDER-DB) [He et al., 2023], NicheNet database [Browaeys et al., 2019], CellChatDB [Jin et al., 2021], and NeuronChatDB [Zhao et al., 2023]. The exFINDER-DB covers a wide range of gene-gene interactions across all three signaling steps, while the NeuronChatDB focuses on neuron-receptor interactions. IncytrDB includes extensive gene-gene interactions for both human and mouse (Supplementary Table 2). Note that IncytrDB may include indirect protein-protein interactions such as those mediated by association or co-expression within the same pathway. For ligand-receptor interaction, however, we require direct interaction between the proteins.

Kinase-substrate interactions predicted by the Kinase Library were appended to IncytrDB for some analyses. If the substrate was in the unmodified DB as a receptor, the Receptor-to-EM database was appended with a substrate-to-kinase interaction. If the substrate was in the unmodified DB as an EM, the Receptor-to-EM database was appended with a kinase-to-substrate interaction, and the EM-to-Target database was appended with a substrate-to-kinase interaction. If the substrate was in the unmodified DB as an EM, the EM-to-Target database was appended with a kinase-to-substrate interaction.

### Ligand-Target Signaling Pathway Inference

The required inputs for the Incytr analysis include user-assigned sender and receiver cell groups (i.e. cell clusters, which may include multiple cell types), and their corresponding gene lists which will be used to infer the ligand-target signaling pathways. These input genes can be derived from prior analysis (e.g., differentially expressed genes) or user-selected. Incytr then uses these lists and the IncytrDB to infer ligand-target pathways where the ligand→receptor (L-R), receptor→effector molecule (R-EM), and effector molecule→target (EM-T) interactions are all documented. More specifically, Incytr constructs the L-R-EM-T signaling pathways (L-T for short) from the IncytrDB which satisfy the following conditions: (1) all ligands are the input sender genes, representing the signals from the sender cell group; (2) the rest of the components are the receiver genes, which are in the receiver cell group; (3) for each L-R-EM-T signaling pathway, its receptor, effector molecule, and target are different. All the inferred pathways are stored as a table, with each row being one L-R-EM-T pathway, and the entire table representing a collection of the ligand-target signaling pathways. In addition, users can filter the database to select only the pathways with their genes of interest by inputting lists of ligands, receptors, EMs, or targets.

### Bulk and Single-cell Data Processing

Single-cell (or nuclei) transcriptomics data is required to perform Incytr analysis, while the proteomics and phospho-proteomics data can be either bulk or single-cell level. Users should ensure that their data has been normalized, scaled, and, if necessary, batch-corrected. Input data will be processed to obtain the cluster-level average expressions.

#### Proteomics and Phospho-proteomics Data Debulking

TMT channels were normalized within each plex. Technical replicates were averaged. Aggregate gene transcript expressions across each sample or experimental group and cell cluster were extracted from the Seurat objects used for analysis. If the transcript raw count for a gene in one sample and cell type was zero, a value of 1 was imputed to resolve later division-by-zero errors when calculating fold changes in signal probabilities between groups (prior to the implementation of imputation, many of the paths with high signal probability fold changes had low probability in one condition and a probability of zero in the other, but the real difference between conditions is small in these cases).

Within each gene and sample, the proportion of the transcript signal coming from each cell type was calculated. For every gene and each sample in the mass spectrometry data, the protein expression signal was proportionally assigned to cell types according to the distribution of the transcript signal in the aggregate single-cell data. Analogously, for phosphorylation data, the phosphoprotein expression signal for a site was proportionally assigned to cell types according to the distribution of the transcript signal from the whole gene in the aggregate single-cell transcriptomic data.

#### Calculation of the Average Expression Levels in Cell Groups

The average expression level of a gene in the cell group *i* is calculated using the normalized single-cell transcriptomics data, which can be obtained by applying the *NormalizeData* function to the count matrix in *Seurat*. Meanwhile, considering the noise effect of the data, a statistically robust mean method has been used [Jin et al., 2021].

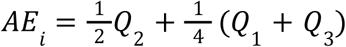

Here *AE_i_* represents the calculated average expression level of the gene, *Q*_1_, *Q*_2_, and *Q_3_* is the first, second, and third quartile of the expression levels of the gene in the corresponding cell group.

#### Integration of the Multi-modal Data

In addition to the single-cell transcriptomics data, Incytr can integrate other multi-modal data (such as proteomics and phosphorylation data) into the analysis. Currently, Incytr allows inputting one proteomics dataset and up to two phosphorylation datasets (such as phospho-S|T and phospho-Y) for analysis. Each set of data should include two separate tables representing the average expression of the genes (row) in every cell group (column) in different conditions. All input datasets should be positive values to avoid possible later division-by-zero errors when performing the cross-condition comparison.

### Quantitative Analysis of the Inferred L-T Pathways

*Incytr performs a series of quantitative and comparative analyses on the inferred ligand-target signaling pathways (from the sender to the receiver cell groups) using single-cell transcriptomic and multi-modal data, which include the following steps:*

#### Prediction of the Signaling Probability

Incytr predicts the signaling probability of each L-T pathway based on the expression levels of the four involved genes using the single-cell transcriptomics data. A Hill function model is used here to predict the interaction probabilities of the following three steps: ligand-receptor, receptor-EM, and EM-target [He et al., 2023].

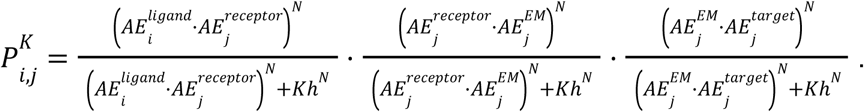

Here, 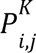 is the signaling probability of the pathway *K* from cell group *i* to *j*, while *K*ℎ is the half-saturation constant (*K*ℎ = 0. 5 by default), and *N* is the Hill coefficient (*N* = 2 by default). It should be noted that changing the values *K*ℎ does not affect relative values between different pathways [He et al., 2023] (Supplementary Figure 9A), so users may adjust their values to have clearer visualizations.

#### Quantification of the Cross-condition Change via the Adjusted Fold Change (aFC) Value

The log2 fold change value is widely used to indicate the difference between two values [Tusher et al., 2001]. However, although two very small numbers may result in an enormous log2-fold change (such as 10^-4 and 10^-6), such a change may not be as biologically significant as the fold change between larger numbers (like 50 and 0.5). Here, Incytr uses the adjusted log2 fold change value (aFC value) which can avoid such false-significant change. If we regard any value below the threshold *a* as non-significant (for example, the 75th percentile o the data), then the aFC value of two numbers *x* and *y* is computed based on the following model:

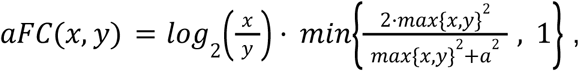

where *aFC* ∈ (− ∞, + ∞). This model ensures that if *x* < *a* and *y* < *a* then *aFC*(*x*, *y*) < *log_2_* (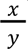) hold, which means when *x* and *y* are both very small, their log2-fold change value will be adjusted to a lower value. The aFC value is used on both single-cell transcriptomics and multi-modal data to quantify the cross-condition change of individual L-T pathways and genes.

#### Identification of the Statistically Significant L-T Pathways

To quantify the statistical significance of a L-T pathway from cell group *i* to *j*, we first performed a permutation test by randomly permuting the labels of all cell groups, re-calculating its communication probability *P_i,j_*. The corresponding *p*-value estimate of each pathway is computed by:

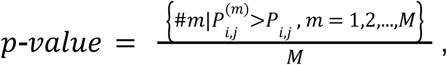

where 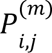 is the signaling probability based on the *m*-th permutation, and *M* is the total number of permutations performed (here *M*=100 by default). The quantification of the statistical significance is based on the single-cell transcriptomics data. The p-value for every pathway is then adjusted for multiple testing (FDR) [Benjamini & Hochberg, 1995]

using the *p.adjust* function from the R package *stats* (the default correction method is “BH”). L-T pathways with adjusted p-values under 0.05 are considered true pathways.

#### Quantification of the Expression Exclusiveness between Cell Groups

The exclusive high expression of a gene in one cell group can indicate some active biological process. For example, a kinase substrate’s exclusive high expression in one cell group can indicate such a group is undergoing a strong kinase-specific phosphorylation. To quantify the expression exclusiveness based on the single-cell transcriptomics data in one condition, we denote *C* to be the set of all cluster labels and define the exclusiveness index (EI) of the gene *g* in cell group *j* (*j* ∈ *C*) in the following way:

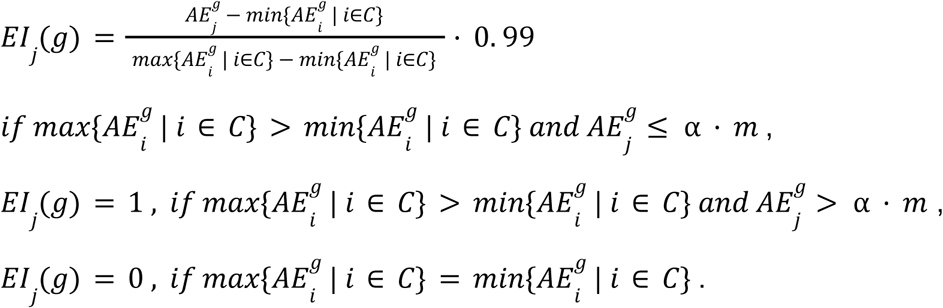

Here 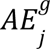 is the average expression of the gene *g* in the cell group *j*, 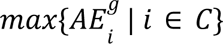 and 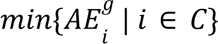 are the highest and lowest average expression levels of *g* in all cell groups, respectively. *m* represents the second highest average expression level, and α is the fold change threshold to determine the “highest exclusive level” (α = 10 by default). Overall, the *EI_j_* comes with the following properties: (1) in general, a higher rank of the 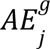 (in all cell groups) will lead to a higher EI; (2) *EI_j_*(*g*) = 1 if and only if *g* has the highest expression level in the group *j* and it is α times higher than the second highest value; (3) *EI_j_*(*g*) = 0 if *g* has the same expression level in all cell groups.

### Kinase Analysis Using the Single-cell Transcriptomics Data

Using the Kinase Library, we can identify the kinases potentially responsible for phosphorylating the proteins within the ligand-target signaling pathways.

#### Identification of the Signaling-involved and Signaling-related Kinases

Kinases inferred to phosphorylation substrates (either R, EM, or T) within these pathways but are not themselves part of the pathways are classified as signaling-related kinases (SrKs) (for example, molecule X from Supplementary Figure 1). Conversely, proteins encoded by genes within the ligand-target signaling pathways could themselves function as kinases, phosphorylating their downstream targets. For example, an EM might serve as a kinase, with the T as its substrate. These kinases, as integral components of the ligand-target pathway, are termed signaling-involved kinases (SiKs) (for example, molecule T2 from Supplementary Figure 1). Once SrKs and SiKs are identified, their relative phosphorylation activity can be quantified in the corresponding cell group using the Exclusiveness Index (EI).

#### Evaluation of the Kinase Analysis Results for Each L-T Pathway

For all SiKs and SrKs, Incytr first calculates their EIs in the receiver cell group. Then Incytr will record the SrKs with the highest EI for future exploration, although these results will not be used in the L-T pathway evaluation module. Moreover, for each L-T pathway, there can be at most six “kinase-substrate” pairs (receptor - EM, receptor-target, EM - target, EM-receptor, target-receptor, or target - EM). To evaluate how the kinase information supports the L-T pathway, Incytr calculates the SiK-score, which is the mean of the kinases’ EI of all six pairs:

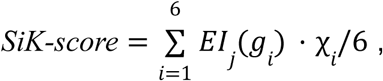

here χ*_i_* = 1 if the corresponding kinase is a SiK, otherwise χ*_i_* = 0. *i* = 1, 2, …, 6 representing the six “kinase-substrate” pairs, and *g* is the kinase in the corresponding pair. Naturally, more identified SiKs with higher EI will lead to a higher SiK-score, indicating a more active phosphorylation process.

#### Quantification of the Signaling Pathway Difference Between Conditions

Incytr provides a comprehensive quantitative evaluation of all L-T pathways on their cross-condition difference based on the overall analysis results, which include the following four parts: (1) the transcriptomics-based pathway differential score (T-PDS) based on the comparative analysis using the single-cell transcriptomics data; (2) the proteomics and phosphorylation-based pathway differential score (P-PDS and Ph-PDS) based on the comparative analysis using the proteomics and phosphorylation data, respectively; (3) the kinase-based pathway differential score (K-PDS) based on the kinase analysis; (4) the pathway differential score (PDS) reporting the overall strength of how much a L-T pathway differs between conditions.

#### Calculation of the Transcriptomics-based Pathway Differential Score (T-PDS)

The T-PDS of an L-T pathway is calculated based on its aFC value on the signaling probabilities between conditions. For a L-T pathway *K*, signaling from the cell group *i* to *j*, its T-PDS is defined based on the following logistic model:

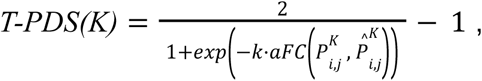

where 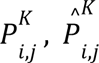 are the predicted signaling probabilities of two conditions, respectively. Here *k* is a positive parameter controlling the sensitivity of T-PDS to the aFC and *k* = 2 by default, noting that changing the values of *k* does not affect the ranking between different L-T pathways (Supplementary Figure 9B). Thus, *T-PDS(K)* ranges from -1 to 1 with a sigmoidal shape, and a higher absolute value indicates a stronger differential activation between the two conditions.

#### Calculation of the Proteomics and Phosphorylation-based Pathway Differential Score (P-PDS and Ph-PDS)

The P-PDS and Ph-PDS of an L-T pathway are calculated based on its components’ aFC values for the proteomics and phosphorylation levels between conditions, respectively. Here we use *m* = 1, 2, 3, 4 to represent different signaling components (ligand, receptor, EM, target). Then the P-PDS and Ph-PDS of the L-T pathway *K* signaling from cell group *i* to *j* are calculated using the same formula:

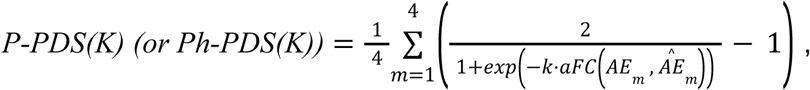

where *AE_m_*, 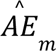 are the average levels of either protein or phospho-protein of the component *m* in its corresponding cell group. Here *k* is the same parameter previously defined. If the input proteomics or phosphorylation data is on bulk resolution, then *AE* represents the average expression level calculated from the debulked data. Noting that for multiple phosphorylation input datasets (such as the data of the phospho-serine and phospho-tyrosine sites), the Ph-PDS of each dataset should be calculated separately. Here, both P-PDS and Ph-PDS range from -1 to 1, showing the overall regulating strength by considering all four components.

#### Calculation of the Kinase-based Pathway Differential Score (K-PDS) and the Pathway Differential Score (PDS)

To calculate the K-PDS of the L-T pathway *K*, its multi-modal differential score (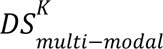) is computed first based on the following model:

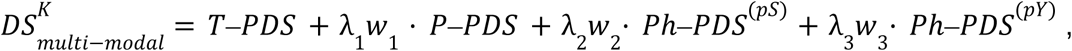

where λ*_i_* = 1 if the corresponding input data is available (proteomics, Ser-Thr phospho-proteomics, Tyr phospho-proteomics, respectively *i* = 1, 2, 3), otherwise λ*_i_* = 0 (*i* = 1, 2, 3). *w_i_* (*i* = 1, 2, 3) represent the weights (*w_i_* = 0. 5, *i* = 1, 2, 3 by default). This multi-modal differential score reflects the differential strength between conditions by considering the added value provided by the multi-modal data. Note that opposite results from different data will counteract the score, ensuring only the signaling pathways supported by the multi-modal data will stand out. Alternate weights can be provided by users who wish to prioritize modalities unequally or exclude any modality. For a L-T pathway *K*, signaling from cell group *i* to *j*, we denote its SiK-score of two conditions as *SiK-score_1* and *SiK-score_2,* respectively. Then, its K-PDS is computed in the following way:

1. if 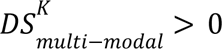 (meaning subpathway *K* is more prominent in condition1), the *K*⎼*PDS* = *SiK*⎼*score* 1;
2. if 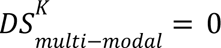, then *K*⎼*PDS* = (*SiK*⎼*score* 1 + *SiK*⎼*score* 2)/2;
3. if 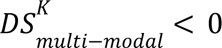, then *K*⎼*PDS* = *SiK*⎼*score* 2.

And its PDS is: *PDS* = 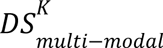 + *w*_4_ · *K–PDS*. Here *w*_4_ is the weight and *w*_4_ = 0. 5 by default.

### Output Visualization

We developed an interactive visualization tool, Incytr-Viz, in order to view, filter, and explore the sets of four-step signaling pathways output by the Incytr analysis package. The tool provides two main interactive graphics (Figure 1E): a cellular interaction graph, which displays the number of pathways between pairs of cell populations, and a river (Sankey) plot, which displays the genes that comprise the pathways. Graphics for each experimental condition/group are separated into individual panels, to allow comparison of predicted pathway sets across conditions. Visualized pathways can also be filtered or highlighted using pertinent Incytr output metrics, such as sender and receiver cell populations, component genes, signalling probability, T-PDS, P-PDS, and, optionally, by user-provided UMAP coordinates.

## Data Availability

The dataset for naïve CD4+ T cell stimulation with IL-6 was obtained from [Ariss et al., 2024].

The A549 Mix-Seq data was sourced from figshare (https://figshare.com/s/139f64b495dea9d88c70), where counts are for all genes unlike the filtered data in PerturBase [McFarland and Paolella et al., 2020].

MC38 dataset: scRNA-seq and processed proteomics data is available on Zenodo (https://zenodo.org/records/14926201).

5XFAD dataset: Processed proteomics data is available on Zenodo (https://zenodo.org/records/14926201). Previously published sequencing data is available in the Gene Expression Omnibus (GEO) database managed by the National Center for Biotechnology Information (NCBI), accession number: GSE140511.

COVID dataset: Previously published sequencing data from the first cohort of HC and COVID-19 [Eddins et al., 2023] is deposited in the Gene Expression Omnibus (GEO) database managed by the National Center for Biotechnology Information (NCBI; accession number: GSE186267). Sequencing and protein expression data for a second cohort are available on Zenodo (https://zenodo.org/records/14926201).

## Code Availability

Incytr-Analysis source code is available at https://github.com/ChanghanGitHub/Incytr Source code for the visualization package (Incytr Viz) is available at https://github.com/cellsignal/incytr-viz

## Acknowledgements

This study was supported in part by the National Institutes of Health (NIH) National Institute of Allergy and Infectious Diseases (NIAID) R01AI123126 (EEBG), R01AI123126-05S1 (EEBG), R21AI167032 (to EEBG), and the Lowance Center for Human Immunology (EEBG). QN was supported by the National Institutes of Health (NIH) grant R01GM152494. DJE was partially supported by Emory’s Laney Graduate School Fellowship.

We acknowledge the use of OpenAI’s ChatGPT for assistance in refining the language and style of this manuscript. The model was utilized to improve clarity and coherence while ensuring that the original scientific content remained intact. All intellectual contributions and interpretations are the sole responsibility of the authors.

## Contributions

Conceptualization and study design: CH, CS, IC, IG, KR, QN, DO;

Data acquisition and curation: BZ, DJE, AK, SS, TL, AP, LH, MA, AFG, DD, AA, EEBG, KR;

Methodology development: CH, CS, IC, ET, IG, KR, QN, DO;

Data analysis and interpretation: CH, CS, BZ, ST, DJE, AK, JY, SS, TL, AP, SHL, DG, AFG, LH, ET, IG, MA, DD, NRR, EEBG, MC, KR, DO;

Manuscript drafting: CH, CS, IC, BZ, ST, DJE, AK, TL, ET, SHL, DG, AFG, IG, MA, DD, NRR, MC, KR, QN, DO;

Critical review and editing: CH, CS, IC, BZ, ST, DJE, AK, JY, SS, TL, AP, LH, ET, SHL, DG, AFG, IG, MA, DD, AA, NRR, EEBG, MC, KR, QN, DO;

Supervision and funding acquisition: KR, QN, DO.

