## Supplementary Materials for "Cell signaling pathways discovery from multi-modal data"

**This PDF file includes:**

Materials and Methods
Supplementary Figures 1 to 9
Supplementary Tables 1 to 3
Captions for Supplementary Data 1 to 3

**Other Supplementary Materials for this manuscript include the following:**

Supplementary Data 1. Benchmarking table comparing the ability of GSEA, exFINDER, and Incytr to rediscover known cancer-specific, 5XFAD model-specific, and COVID-specific pathways.

Supplementary Data 2. A list of references supporting the primary pairwise L-R, R-EM, and EM-T interactions presented in this paper (Figures 3-5).

Supplementary Data 3. Output of the Incytr analysis on the 5XFAD dataset, focusing on interactions involving Apoe, App, Trem2, and Lrp1.

### Materials and Methods

#### Data/Sample Overview and Analysis

##### Method Validation Using Small-Molecule Perturbation Assay Data from a Lung Cancer Cell Line

###### Mix-Seq Data Processing

Single-cell RNA-sequencing data were obtained from Mix-Seq Experiment 10, a multiplexed drug perturbation experiment [McFarland et al., 2020]. Data were sourced from figshare accession 10298696. Cell line identity labels (singlet_ID) were obtained from per-drug classifications CSV files provided in the figshare repository. The concatenated table was merged with the AnnData observation metadata to retain only barcodes present in both the PerturBase-filtered expression matrix and the classifications table. Cells were filtered to retain only those classified as `cell_quality == 'normal'`, and cells with fewer than 200 total counts and genes detected in fewer than 10 cells were removed. This resulted in 20–102 cells per condition.

###### Mix-Seq Incytr Analysis and MSigDB Gene Set Mapping

The top 6,000 genes were selected by ranking mean expression computed via Seurat's `AverageExpression` function across all cells using the raw counts layer. L-R-EM-T chain analysis was performed using Incytr. For each drug perturbation, A549_LUNG cells were designated as both the sender and receiver population, and a two-condition comparison was run between vehicle control (CTRL) and the drug-treated condition. Incytr hits were annotated against the MSigDB C2 curated gene set collection (c2.all.v2025.1.Hs.symbols.gmt). Per-drug pathway counts were computed by tallying the number of significant L-R-EM-T chains mapping to each pathway term. Downstream analyses were restricted to pathways with SigProb > 0 and aFC values either < −1 or > 1. A negative aFC value (aFC<-1) means the pathway is upregulated in drug treatment, whereas aFC>1 represents downregulated pathways with the drug treatment. Row-wise z-score normalization was performed, followed by hierarchical clustering of pathways using Euclidean distance and Ward linkage.

#### Method Validation Using the Mouse Immune Data

##### Data Processing and Incytr Analysis

The scRNA-seq data of the cytokine-treated and PBS-treated cells of 14 cell types was obtained from the previous study of Cui et al. [Cui et al., 2024], and 14 cytokines were selected for method validation based on the cytokine–cell type combinations reported in the original study (Supplementary Figure 1 of [Cui et al., 2024]). For each cytokine, the following Incytr analysis between the control and perturbed conditions was performed within a given cell type: (1) PBS-treated cells were used as both sender and receiver cells under the control condition; (2) Cytokine-treated cells were used as the receiver cells under the perturbed condition; (3) To simulate cytokine injection in the sender cells, we used PBS-treated cells as the sender population and manually perturbed the cytokine expression to the 85th percentile of its expression distribution; (4) Only the signaling pathways with abd(PDS)>=0.2 were kept, while the permutation test was skipped due to the computation cost.

##### Hallmark ID Mapping of the Inferred Signaling Pathways

Hallmark IDs were identified by mapping or overlaying Incytr networks and cytokine dictionary gene sets onto the Hallmark gene sets from the MSigDB database (https://www.gsea-msigdb.org/gsea/msigdb/), in order to obtain enriched biological states or processes. We required that all four genes representing L, R, EM, and T in an Incytr-derived network map to Hallmark gene sets for a given Hallmark ID to be assigned. For the gene programs listed in Supplementary Table 5 of [Cui et al., 2024], we applied a ≥3 gene threshold, that is, a gene program was mapped to a Hallmark ID if at least three of its genes were found in that Hallmark gene set. This latter condition was applied to cytokine-influenced signaling networks, that is, networks that are affected by the cytokine but do not necessarily include the cytokine molecule directly.

##### Results Validation via the Previous Study

First, the Hallmark Pathway IDs associated with the cytokine treatments reported in the original study [Cui et al., 2024] were obtained from the shared source data and Supplementary Figure 1. Second, for each cytokine within a given cell type, we selected the Hallmark Pathway IDs identified from the cytokine-driven signaling pathways (i.e., where the selected cytokine serves as the ligand), and assessed which of these were consistent with the results reported in the original study. The results are shown in Figure 2A. Thirdly, for the Hallmark Pathway IDs identified from non-cytokine-driven signaling pathways, we only kept those which have at least one pathway gene (i.e., ligand, receptor, EM, or target) included in the gene list that Cui et al. used to identify the Hallmark Pathway IDs. Furthermore, we compared them to more constrained paper-reported results to ensure robustness -- only the paper-reported Hallmark Pathway IDs which have at least three common genes mapped to the Hallmark database were used for comparison. The results are shown in Supplementary Figure 3.

#### MC38 Dataset

##### Generation of the MC38 Mouse Model

###### Cell Culture

The MC38 murine colon adenocarcinoma cell line was obtained from Kerafast and cultured according to standard protocols. Cells were maintained in high-glucose Dulbecco's Modified Eagle Medium (DMEM; Gibco, Thermo Fisher Scientific) supplemented with 10% heat-inactivated fetal bovine serum (FBS; Gibco), 1% penicillin-streptomycin (Pen/Strep; Gibco), and 2 mM L-glutamine at 37°C in a humidified atmosphere containing 5% $CO_2$.

###### Preparation of Cells for Injection

MC38 cells were harvested during the logarithmic growth phase to ensure high viability and proliferation rates. Cells were detached using 0.25% trypsin-EDTA (Gibco) and resuspended in sterile phosphate-buffered saline (PBS; pH 7.4). Cell concentration and viability were assessed using a hemocytometer and trypan blue exclusion method, respectively. Cell suspensions were adjusted to a final concentration of 1 × $10^6$ cells per 100 µL of PBS for injection.

##### Animal Handling and Ethical Approval

All animal procedures were conducted in compliance with the Institutional Animal Care and Use Committee (IACUC) guidelines of Bluefin Biomedicine. Female C57BL/6 mice, aged 6–8 weeks, were purchased from The Jackson Laboratory and housed under specific pathogen-free conditions with a 12-hour light/dark cycle. Mice were allowed to acclimate for one week prior to experimentation and had ad libitum access to food and water.

##### Subcutaneous Tumor Implantation

For tumor induction, mice were briefly anesthetized with isoflurane inhalation (induction at 3–4%, maintenance at 1–2%). Using a 27-gauge needle, 0.5 × $10^6$ MC38 cells in 100 µL of PBS were injected subcutaneously into the right flank of each mouse. Mice were monitored until full recovery from anesthesia and then returned to their cages.

##### Tumor Monitoring

Tumor growth was measured three times per week using digital calipers. Tumor volume was calculated using the formula:

*Tumor volume* = 1/2(*length* × *width*^2)

where length is the longest dimension and width is the dimension perpendicular to the length. Mice were also monitored for signs of discomfort, weight loss, or any adverse effects. Humane endpoints were established, and mice were euthanized if tumor volume exceeded 1,500 $mm^3$ or if they exhibited signs of distress.

##### MC38 scRNAseq Data Generation and Analysis

MC38 tumor samples were dissociated using a GentleMACs, and mouse tumor dissociation kit (Miltenyi Biotec). Cell suspensions were subsequently labeled using Biolegend’s TotalSeq-A anti-mouse hashtag antibodies. Single cell suspensions were counted, pooled, and analyzed following 10X Genomics Single Cell 3' v2 assay.

Resulting libraries were analyzed on an Illumina NovaSeq SP 100-cycle run. The resulting sequencing data was analyzed using CellRanger 3.1.0, and searched with mm10-3.0.0 reference.

Seurat was used to perform quality control, normalization, scaling, dimensionality reduction, and clustering. Marker genes were used to annotate cell types. To distinguish between EMT cells and other cell types we relied on previously reported EMT signatures [Taube et al., 2010; Anastassiou et al., 2011; Tian et al., 2020; Gilles et al., 2013]. In particular, Dcn, Serpinh1, Mmp9, Serpinf1, Zeb1 and Col6a2 genes were found to be upregulated in EMT cells (as well as some fibroblasts) and not in cancer cells.

#### MC38 Proteomics Data Generation

##### Lysis and Protein Extraction

Flash frozen samples were resuspended and probe tip sonicated in 8M urea with 200mM HEPEs pH8.5 with 1x Protease and Phosphatase inhibitors (CST #5872). Samples were then reduced for 30 minutes at 50C with 5mM DTT. Samples were alkylated in the dark for 30 minutes at room temperature with 10mM iodoacetamide. Alkylation was quenched with 5 mM DTT.

##### Protein Digestion

Samples were diluted to 2M urea with 20mM HEPEs pH 8.5 with 1mM CaCl2, and digested overnight at 37C with LysC (Wako-Chem). Samples were further diluted to 1M urea with 20 mM HEPEs pH 8.5 with 1mM CaCl2 and digested for 6 hours with Trypsin (Pierce). Samples were acidified with trifluoroacetic acid (TFA) de-salted with a Waters SepPak C18 column. Peptide concentrations were estimated with a microBCA kit from Pierce (#23235). Peptides were TMT-labeled in 200 mM HEPEs pH 8.5 with 30% acetonitrile for 1 hour, combined, dried in a speedvac, and desalted with a SepPak C18 cartridge.

##### Total Protein Analysis Sample Preparation

For total protein analysis, 100 ug of combined peptide material was resuspended in buffer A (10mM NH 4 HCO 2, pH10, 5% ACN), and fractionated on a Zorbax Extended C18 column (2.1 × 150 mm, 3.5 μm, no. 763750-902, Agilent) using a gradient of 10-40% bRP buffer B (10 mM NH 4 HCO 2, pH10, 90% ACN). 96 fractions were collected and concatenated into 24 fractions. Fractions were dried, de-salted with stop-and-go extraction tips (STAGE-tips), and dried for LC-MS analysis.

##### Enrichment of Post-translationally Modified Peptides

3mgs of combined labeled peptides were IMAC enriched using High-Select TM Fe-NTA Phosphopeptide Enrichment Kits (Thermo Fisher Scientific, San Jose, CA). The remainder of the labeled pool of peptides was analyzed using PTMScan® HS Phospho-Tyrosine (P-Tyr-1000) Kit #38572 from Cell Signaling Technologies. The resulting phosphopeptide elution from IMAC enrichment was fractionated as described above, using a gradient from 5-40% Buffer B. 96 fractions were collected and concatenated into 12 fractions. Fractions were dried, de-salted with STAGE-tips, and dried for LC-MS analysis.

##### LC–MS Analysis of Total Protein

Samples were analyzed on an Orbitrap Fusion Lumos mass spectrometer (Thermo Fisher Scientific, San Jose, CA) coupled with a Proxeon EASY-nLC 1200 liquid chromatography (LC) pump (Thermo Fisher Scientific, San Jose, CA). Peptides were separated on a 100 μm inner diameter microcapillary column packed with ~40 cm of Accucore150 resin (2.6 μm, 150 Å, ThermoFisher Scientific, San Jose, CA). For each analysis, we loaded approximately 1 μg onto the column. Peptides were separated using either a 2.5 h gradient of 6–30% acetonitrile in 0.1% formic acid with a flow rate of 550 nL/min. Each analysis used an SPS-MS3-based TMT method [Ting et al., 2011; McAlister et al., 2014] , which has been shown to reduce ion interference compared to MS2 quantification [Paulo et al., 2016]. The scan sequence began with an MS1 spectrum (Orbitrap analysis, resolution 120,000; 350-1400 m/z, automatic gain control (AGC) target $4.0 \times 10^{\wedge 5}$, maximum injection time 50 ms). Precursors for MS2/MS3 analysis were selected using a Top10 method. MS2 analysis consisted of collision-induced dissociation (quadrupole ion trap; AGC 2.0 × 10^$^4$; normalized collision energy (NCE) 35; maximum injection time 22 ms). Following the acquisition of each MS2 spectrum, we collected an MS3 spectrum, a method in which multiple MS2 fragment ions are captured in the MS3 precursor population using isolation waveforms with multiple frequency notches [McAlister et al., 2014]. MS3 precursors were fragmented by HCD and analyzed using the Orbitrap (NCE 65, AGC 3.5 × 10^$^5$, maximum injection time 120 ms, isolation window 1.2 Th, resolution was 50,000 at 200 Th).

##### LC–MS Analysis of PTM Enriched Fractions

For the analysis of PTM-enriched samples, the same MS and HPLC instruments were used as described above. For each analysis, approximately 500 ngs of enriched peptides were loaded on the column and run over a 120-minute gradient of 2-32% acetonitrile in 0.1% formic acid with a flow rate of 400 nL/min. MS1 spectra were collected in the Orbitrap at a resolution of 60,000 with a scan range of 300-1500 m/z using an AGC target of 4.0e5 with a maximum injection time of 25ms. Peptides for MS2 analysis were isolated using the quadrupole with an isolation window of 0.8 m/z. MS2 spectra were generated using Higher-energy collision dissociation (HCD) with a collision energy of 40%. Fragments were collected in the Orbitrap at a resolution of 50,000 with a first mass of 110 m/z, an AGC target of 5.0e4, and a maximum injection time of 200ms.

#### 5XFAD Dataset

##### 5XFAD snRNAseq Data

We used previously published snRNA data from 5XFAD and WT mouse brains (n = 3) [Zhou et al., 2020]. Seurat was used to perform quality control, normalization, scaling, dimensionality reduction, and clustering. Cell types were identified according to the expression of marker genes selected in the original study.

##### 5XFAD Proteomics Data Generation

One male and one female half mouse brains from WT and 5XFAD backgrounds were analyzed. Samples were additional replicates from the study from which the matched snRNAseq data were generated [Zhou et al., 2020]. Total and phosphoproteome datasets were generated and analyzed as described above for the MC38 tumor samples with the following exceptions. After sample lysis, protein concentration was estimated using microBCA kit from Pierce. 1 mg of protein per sample (in triplicate) was then reduced, and alkylated as described above. Proteins were then precipitated using a MeOH Chloroform extraction. Protein pellets were resuspended in 200 uL of 200 mM EPPS buffer (pH 8.5), digested with 10µg of LysC overnight at 37C, and 10 µg of Trypsin was added for an additional 6-hour digestion. Peptides were then TMT labeled in 200mM EPPS with 30% AcN for 1 hour at room temperature using TMT-pro reagents from Thermo.

##### Bulk Total Proteomics Data Analysis

Mass Spectra were processed using a Comet-based software pipeline [Eng et al., 2012; Huttlin et al., 2010]. Resulting data were searched with a fully tryptic database containing mouse Swissprot consensus entries plus isoforms allowing for a static modification of lysine and N-termini with TMT (229.1629 Da) or TMT-pro (304.2071 Da) and carbamidomethylation (57.0215 Da) of cysteine, along with variable oxidation (15.9949 Da) of methionine. Searches were performed using a 20 ppm precursor ion tolerance, the production tolerance was set to 1.0 Th. Peptide-spectrum matches (PSMs) were adjusted to a 2% false discovery rate (FDR) using previously described linear discriminant analysis [Elias & Gygi, 2007; Elias & Gygi, 2009]. Filtered PSMs were collapsed to a final protein-level FDR of < 2%. Protein assembly was guided by principles of parsimony to produce the smallest set of proteins necessary to account for all observed peptides [Huttlin et al., 2010]. MS3 spectra with TMT reporter ion summed signal-to-noise ratios less than 100 were excluded from quantitation [McAlister et al., 2012].

##### Bulk Phosphorylation MS Data Analysis

PTM data was analyzed as described above, with a fragment ion tolerance of 0.02 Th instead. Variable modifications were considered for phosphorylation on serine, threonine, and tyrosine (79.9663 Da). PTM sites with A-score values > 13 were considered as localized for downstream analysis [Huttlin et al., 2010; Beausoleil et al., 2006].

##### Kinase-Substrate Matching

Phosphorylation sites within the provided phosphoproteomic datasets (MC38 and 5XFAD) were analyzed for associated kinases using previously described methods [Johnson et al., 2023]. Briefly, 10-mer segments surrounding S/T/Y residues were compared against experimentally determined sequence motifs representing each kinase's most biochemically favorable substrate and assigned a score. Kinases were then ranked according to the percentile of that site's score with respect to scores for all sites in the phosphoproteome for that kinase. Kinase-substrate matching is performed by determining the (15 for phosphoY, 5 for pS/pT) most highly ranked kinases for a given site. Site scoring is publicly available at https://kinase-library.phosphosite.org/.

#### COVID-19 Dataset

There are two cohorts of healthy control (HC) and COVID-19 patient samples that were used in this study. The first cohort (six HC and four COVID-19 patient samples previously sequenced) and a second cohort (three HC and three COVID-19 patient samples not previously sequenced) collected as part of the previously published report [Eddins et al., 2023] were processed as part of this study. For metadata associated with these patients see Supplemental Table 1 from Eddins et al., 2023 [Eddins et al., 2023]. The first cohort patient IDs: PHA1005, PHA1006, PHA1007, PHA1008, PHA1009, PHA1010, PHA0007, PHA0008, PHA0010, PHA0018; the second cohort patient IDs: PHA1005, PHA1007, PHA1008, PHA0001, PHA0002, PHA0003.

Whole blood samples from consenting donors were collected and processed under institutional review board (IRB)-approved protocols as previously described [Eddins et al., 2023]. Following plasma isolation, red blood cells (RBCs) were depleted from whole blood using the EasySep™ RBC Depletion Reagent (STEMCELL Technologies). Following RBC-depletion, cells for scRNA-seq were surface stained with a panel of 89 antibody-derived tags (ADT; see Supplemental Table 4 in [Eddins et al., 2023]) and loaded to target encapsulation of 10,000 cells using a Chromium Controller (10X Genomics, Pleasanton, CA). Gene expression (GEX) and ADT libraries were generated using the Chromium Single Cell 5′ Library & Gel Bead Kit v1.1 with feature barcoding protocol per the manufacturer's instructions and sequenced in a S4 flow cell on a NovaSeq 6000 (Illumina, San Diego, CA). Peripheral blood mononuclear cells (PBMCs) were isolated using either the EasySep PBMC isolation kit or Lymphoprep (StemCell Technologies), from the remaining cells and were frozen at -196˚C in 10% ultra-pure grade (≥99.9%) dimethyl sulfoxide (DMSO; VWR) in fetal bovine serum (FBS, Thermo Fisher Scientific) for future analyses.

##### InTraSeq Experiment

Frozen PBMCs from Healthy and COVID-19-infected donors from the second cohort were obtained from the Ghosn lab. The cells were washed once with 10 mL of Roswell Park Memorial Institute (RPMI)-1640 medium, then spun down at 300 x g for 5 minutes at 4°C. The cells were washed again with 10 mL of Phosphate-Buffered Saline (PBS), spun down at 300 x g for 5 minutes at 4°C, then were resuspended in 500 uL of PBS prior to be subjected to the InTraSeq protocol (CST #82906) [Ariss et al., 2024]. The Following conjugates were added in the Immunostaining step: InTraSeq™ 3' Conjugate Antibody Cocktail 1 (CST #48167), S100A9 (D5O6O) Rabbit mAb (InTraSeq™ 3' Conjugate 3004) #97056, T-bet/TBX21 (D6N8B) XP® Rabbit mAb (InTraSeq™ 3' Conjugate 3009) #57412, CD68 (D4B9C) XP® Rabbit mAb (InTraSeq™ 3' Conjugate 3012) #29105, CD3 (UCHT1) Mouse mAb (InTraSeq™ 3' Conjugate 3028) #72489, CD4 (RPA-T4) Mouse mAb (InTraSeq™ 3' Conjugate 3029) #87589, CD8α (SK1) Mouse mAb (InTraSeq™ 3' Conjugate 3030) #25292, Mouse (G3A1) mAb IgG1 Isotype Control (InTraSeq™ 3' Conjugate 3001) #59605.

##### 10x Genomics and Illumina Sequencing

Following InTraSeq, the cells in the second cohort of the COVID-19 dataset were processed using the 10x Genomics Next-Gem 3' single cell kits and User Guide CG000317. After the single-cell RNA and protein library preparation was generated, the samples were pooled and sequenced using the NextSeq 2000 and the P2 or P3 kits (100 cycles). Base-calling was performed using NextSeq 1000/2000 Control Software v1.5.0 (RTA v3.10.30).

##### Single Cell InTraSeq Analysis

InTraSeq data were loaded into a Seurat object along with the RNAseq data. Cells were filtered on the number of features (500-3000) and counts (900-7000) in the RNA data, the number of counts (<10000) in the protein data, and the percentage of mitochondrial genes (<5%). The protein data was normalized using centered log ratio transformation. Proteins were visualized using a minimum cutoff of quantile5 or quantile10 and a maximum cutoff of quantile90 or quantile95 expression.

#### HIV Dataset

Archived PBMC samples were previously obtained from 8 individuals living with HIV who were virally-suppressed and participating in the UCSF SCOPE cohort. PBMCs from each donor were thawed at 37 °C, washed once with warm media (RPMI (Corning Inc., Corning, NY) supplemented with 10% FBS (VWR, Radnor, PA)), and then resuspended in FACS buffer (RPMI, supplemented with 2% FBS and 2 mM EDTA (Thermo Fisher Scientific)). Half of each PBMC specimen was reserved for subsequent CyTOF analysis (refer to section below), while lymphocytes were enriched from the remaining cells through negative selection using the EasySep Direct Human Total Lymphocyte Isolation Kit (StemCell).

##### snRNAseq and snATACseq

###### Cell preparation for simultaneous snRNA and snATAC sequencing

Nuclei were isolated from PBMCs according to the 10X Genomics' (Pleasanton, CA) Demonstrated Protocol CG000365: Nuclei Isolation for Single Cell Multiome ATAC + Gene Expression Sequencing. Briefly, cells were washed twice with cold PBS supplemented with 0.04% BSA, were passed through a 40 μm strainer, and were counted. A maximum of 1 million cells were resuspended in 100 μl of cold Lysis Buffer (10 mM Tris-HCl at pH 7.4, 10 mM NaCl, 3 mM MgCl2, 0.1% Tween-20, 0.1% NP-40, 0.01% digitonin, 1% BSA, 1 mM DTT, 1 U/mL RNase inhibitors, all in nuclease-free water) and incubated for 3 minutes on ice. Cell lysis reaction was quenched with the addition of 1 mL of Wash Buffer (10 mM Tris-HCl at pH 7.4, 10 mM NaCl, 3 mM MgCl2, 1% BSA, 0.1% Tween-20, 1 mM DTT, and 1 U/mL RNase inhibitors in nuclease-free water). Cells were centrifuged for 5 minutes at 4 °C to repellet, followed by two more washes. Cells were then suspended in diluted Nuclei Buffer (provided at 20x concentration by 10X Genomics in the Chromium Next GEM Single Cell Multiome ATAC Kit A, PN-1000280) supplemented with 1 mM DTT and 1 U/mL RNase inhibitors. GEM generation and cell barcoding were immediately performed using the 10X Genomics' Chromium Controller and 10X Genomics' Next GEM Chip J. The construction of ATAC and gene expression libraries was performed in accordance with the 10X Genomics' Next GEM Single Cell Multiome ATAC + Gene Expression User Guide (CG000338, Rev E). Sequencing was performed on the Illumina NovaSeq platform at the UCSF CAT Core facility, following the manufacturer's instructions.

##### Processing snRNAseq Data

Demultiplexed fastq files were aligned to the hg38 transcriptome reference using the 10X Genomics' Cell Ranger v7.0.0 count pipeline [Zheng et al., 2017], with the inclusion of introns enabled. The count matrices were filtered to retain cells with less than 20% percent mitochondrial gene content, a minimum library size of 200 genes, and the top 5% of library sizes were excluded to mitigate the presence of multiple cells within a droplet. The samples were then integrated using Seurat v4.3.1 [Hao et al., 2021]. Biological and technical variables were visualized through Seurat's DimPlot function to evaluate the presence of batch effects in the UMAPs.

##### Annotating Cell Types Using snRNA Data

Cells were mapped with a PBMC reference dataset [Hao et al., 2021] using Azimuth (Butler et al., nd; https://github.com/satijalab/azimuth). Cells exhibiting discordant subtype annotations and low prediction scores (subtype prediction score of < 0.5) were excluded from further analyses.

##### Processing snATACseq Data

The snATAC data was processed in conjunction with the snRNA data utilizing the same Cell Ranger command previously mentioned. The fragments files were then imported into ArchR v1.0.2 [Granja et al., 2021], where cells were filtered on a minimum TSS score of 10 and a minimum of 1000 fragments using the createArrowFiles function. Cell types and subtypes were assigned by aligning cell barcoded with the snRNA annotations.

#### CyTOF

##### Preparation of samples for CyTOF

A total of 0.5-6 million cells per sample were treated with cisplatin (Sigma-Aldrich) as a live/dead indicator, and then fixed with paraformaldehyde (PFA) as previously described [George et al., 2025; Ma et al., 2026; George et al., 2022]. Briefly, cisplatin staining was performed by incubating cells for 60 sec in 4 ml contaminant-free PBS (Rockland) containing 2 mM EDTA (Corning) and 12.5 μM cisplatin (Sigma-Aldrich). Cisplatin staining was then quenched with 10 ml of CyFACS (contaminant-free PBS (Rockland) supplemented with 0.1% bovine serum albumin (BSA; Sigma-Aldrich) and 0.1% sodium azide (Sigma-Aldrich)). The cells were then pelleted by centrifugation and fixed for 10 minutes at room temperature (RT) in 2% PFA diluted in CyFACS. Cells were then washed twice with CyFACS, resuspended in 10% DMSO in CyFACS, and stored at -80°C until CyTOF analysis.

##### CyTOF antibody conjugation, staining, and data acquisition

Our CyTOF panel (Supplementary Table 3) consisted of markers to define T and NK cells (e.g. CD3, CD14, CD4, and CD8) and antiviral responses (e.g. MX1, MX2, AIM2, IFIT3, and pIRF3). Of note, these antigens have been previously validated and analyzed in the context of HIV [George et al., 2025]. Briefly, CyTOF antibodies were conjugated using the Maxpar X8 Antibody Labeling Kit (Standard BioTools) according to manufacturer’s instructions. Briefly, 5 μl lanthanide solution was combined with 0.1 mg Maxpar Polymer Reagent dissolved in 95 μl L-buffer. After incubating for 60 min at RT, 200 μl C-buffer was added to the polymer mixture, transferred into a 3-kD AmiconTM Ultra tube (Fisher), and then centrifuged for 12,000 g for 25 min at RT. The remaining polymer mixture was then washed with 400 μl C-buffer, centrifuged again, and the flowthrough discarded. Concurrently, 100 μg antibody was mixed with 300 μl R-buffer and transferred to a 50-kD AmiconTM Ultra tube (Fisher), which was then centrifuged at 12,000 g for 10 min at RT. A total of 100 μl TCEP (Pierce) diluted in R-buffer (4 mM final concentration) was added to the buffer-exchanged antibody retentate on the column, followed by vortexing and incubation in a 37°C water bath for 30 min. The antibody mix was then washed with 300 μl C-buffer, centrifuged at 12,000 g for 10 min at RT, and the flowthrough discarded. The antibody mix was washed a second time with 400 μl C-buffer and centrifuged at 12,000 g for 10 min at RT. The column was then transferred to a new collection tube. The polymer mixture was then resuspended with 200 μl C-buffer, added to the antibody-polymer mix, and incubated for 90 min. The coupled antibody-polymer mix was then washed with 300 μl W-buffer and then centrifuged at 12,000 g for 10 min at RT. The antibody-polymer mix was then washed three more times with 400 μl W-buffer, and resuspended in 100 μl W-buffer. Centrifugation of the column upside-down at 1,000 g for 2 min in a new collection tube was used to recover the conjugated CyTOF antibody. Conjugated CyTOF antibody was quantified by

NanoDrop (Thermo Fisher), mixed with 100 μl of Antibody Stabilizer (Candor Biosciences), and stored at 4°C.

CyTOF barcoding and staining was conducted similar to methods previously implemented [George et al., 2025; Ma et al., 2026; George et al., 2022]. Briefly, cisplatin-treated samples were thawed and barcoded using Cell-ID 20-Plex Pd Barcoding Kit (Standard BioTools) per manufacturer's instructions. Barcoded samples were then pooled into a single sample and diluted to a concentration of 6 million cells / 200 μl CyFACS per well in Nunc 96 DeepWell polystyrene plates (Thermo Fisher). Cells were blocked with 3 μl of mouse (Thermo Fisher), 3 μl of rat (Thermo Fisher), and 0.6 μl of human AB (Sigma-Aldrich) sera in 200 μl of CyFACS for 15 minutes at 4°C and then washed twice with CyFACS. Cells were then stained with surface CyTOF antibodies (Supplementary Table 3) for 45 min at 4°C, washed twice with CyFACS, and fixed overnight in 100 μl of 2% PFA in CyPBS (metal contaminant-free PBS (Rockland) supplemented with 1 ml 0.5 M EDTA (Corning)). The following day, cells were permeabilized in 200 μl Foxp3 Intracellular Fixation & Permeabilization Buffer (eBioscience) for 30 min at 4°C. Cells were washed twice with Permeabilization Buffer (eBioscience) and blocked in 75 μl Permeabilization Buffer containing 15 μl of mouse and 15 μl of rat sera for 15 min at 4°C. After another two washes with Permeabilization Buffer, cells were stained with intracellular CyTOF antibodies (Supplementary Table 3) for 45 min at 4°C. After an additional CyFACS wash, cells were stained for 20 min at RT with 250 nM of Cell-ID Intercalator-IR (Standard Biotools). After two more washes with CyFACS, cells were fixed overnight in 100 μl of 2% PFA in CyPBS.

Prior to sample acquisition, cells were washed once with Cell Staining Buffer (CSB; Standard Biotools), once with Cell Acquisition Solution buffer (CAS; Standard Biotools), and resuspended in CAS buffer containing 10% (v/v) EQ Four Element Calibration Beads (Standard Biotools). Data were acquired on a Helios CyTOF instrument at the UCSF Parnassus Flow Core. A sample pressure between 4 and 6, and a running speed of 250 to 400 events per second was maintained during acquisition to reduce clogging. Raw data were converted into flow cytometry standard (.fcs) files for further analyses.

Data processing and normalization

CyTOF datasets were concatenated, normalized to EQ calibration beads, and de-barcoded, according to the manufacturer's instructions (Standard Biotools). Identical anchor samples aliquoted from the same PBMC donor were included in each CyTOF run were used to normalize batches of experimental datasets using the application CUHIMSR/CytofBatchAdjust in R [Schuyler et al., 2019]. FlowJo software (version 10.9.0, BD Biosciences) was used to generate 2D plots and identify events corresponding to intact cells (191Ir+193Ir+), live cells (195Pt-), and singlet cells (based on event length). Intact, live, singlet cells were sub-gated into CD8+ T cells, by sequentially gating on CD3+CD14- T cells and CD4-CD8+ T cells, and NK cells, by sequentially gating on CD14- cells, CD3-CD4- cells, and CD7+ cells.

#### Mapping Incytr-identified Pathways onto WikiPathways

To interrogate and gain more insight into the biological significance of all the paths predicted by Incytr, we downloaded and mapped these four-step paths onto WikiPathways gene sets [Agrawal et al., 2024]. These gene sets are derived from the WikiPathways biological pathway database (https://www.wikipathways.org/) and downloaded from the Molecular Signatures Database (MSigDB) (https://www.gsea-msigdb.org/gsea/msigdb) [Liberzon et al., 2011] curated gene set collections, specifically the canonical pathway subcollections. There are 830 human and 189 mouse gene sets/pathways, which are canonical representations of signaling pathways compiled by domain experts. We developed a Python script that overlays the four components of Incytr paths onto WikiPathways. To extend the coverage, we converted mouse gene names to their human equivalents. The Python script is provided and can be downloaded from our software package.

#### Liceptor, Biomarker, and Protein Degraders Databases

The Liceptor database (https://www.evolvus.com/data/liceptor-database) is a gold standard in small-molecule drug discovery. With over 10 million compounds, it is the world's most comprehensive and largest small-molecule ligand bioactivity dataset built by extracting and curating data from patents drafted in English, Japanese, Chinese, and Korean. It includes 2D structures, associated molecular descriptors, and bioactivity data such as assays, functions, and therapeutic indications. The database focuses on various target families - GPCRs, ion channels, transporters, kinases, proteases, phosphatases, nuclear receptors, and cytochrome P450 covering over 19000 targets.

The Biomarker database (https://www.evolvus.com/data/biomarker-database) is a database of Oncology Biomarkers, a curated database of genomic/molecular alterations, investigational and approved therapies and clinical trials. This curated database includes, the various biological, chemical and clinical data points, including but not limited to other properties of the therapies against the biomarkers, if present. The information is derived from different sources such as – trial registries, regulatory filings, labels.

Proximers - The Protein Degraders database (https://evolvus.com/data/proximers-database) is the world's largest curated resource that compiles information on bi-functional and hetero bi-functional compounds, along with their associated bioactivity and pharmacokinetic metadata. The data is sourced from various sources, including patents drafted in English and non-English languages.

The database includes chemical and biological data points related to the following components: protein binding moiety, ubiquitin ligase binding moiety, linker, protein degrader.

#### Incytr-discovered Pathway Embeddings

An aggregation of Incytr-derived four-step signaling pathways was used to train the Doc2Vec model (https://radimrehurek.com/gensim/models/doc2vec.html), independently for each dataset (~1M pathways for MC38, ~96k for 5XFAD, and ~116k for COVID) with the following input parameters: vector_size=60, window=2, min_count=1, epochs=10. All gene names were converted to lowercase to prevent artificial heterogeneity between mouse and human genes, as human gene names are typically written in all capital letters. After training, the embeddings were extracted and concatenated with the quantitative information related to the adjusted log2 fold change value (aFC value) calculated for each pathway using signaling probabilities $P_{i,j}^{k}$ between conditions (e.g., 5XFAD vs WT). These concatenated data were visualized using UMAP (https://umap-learn.readthedocs.io/en/latest/), with default settings (n_neighbors=5, random_state=42).

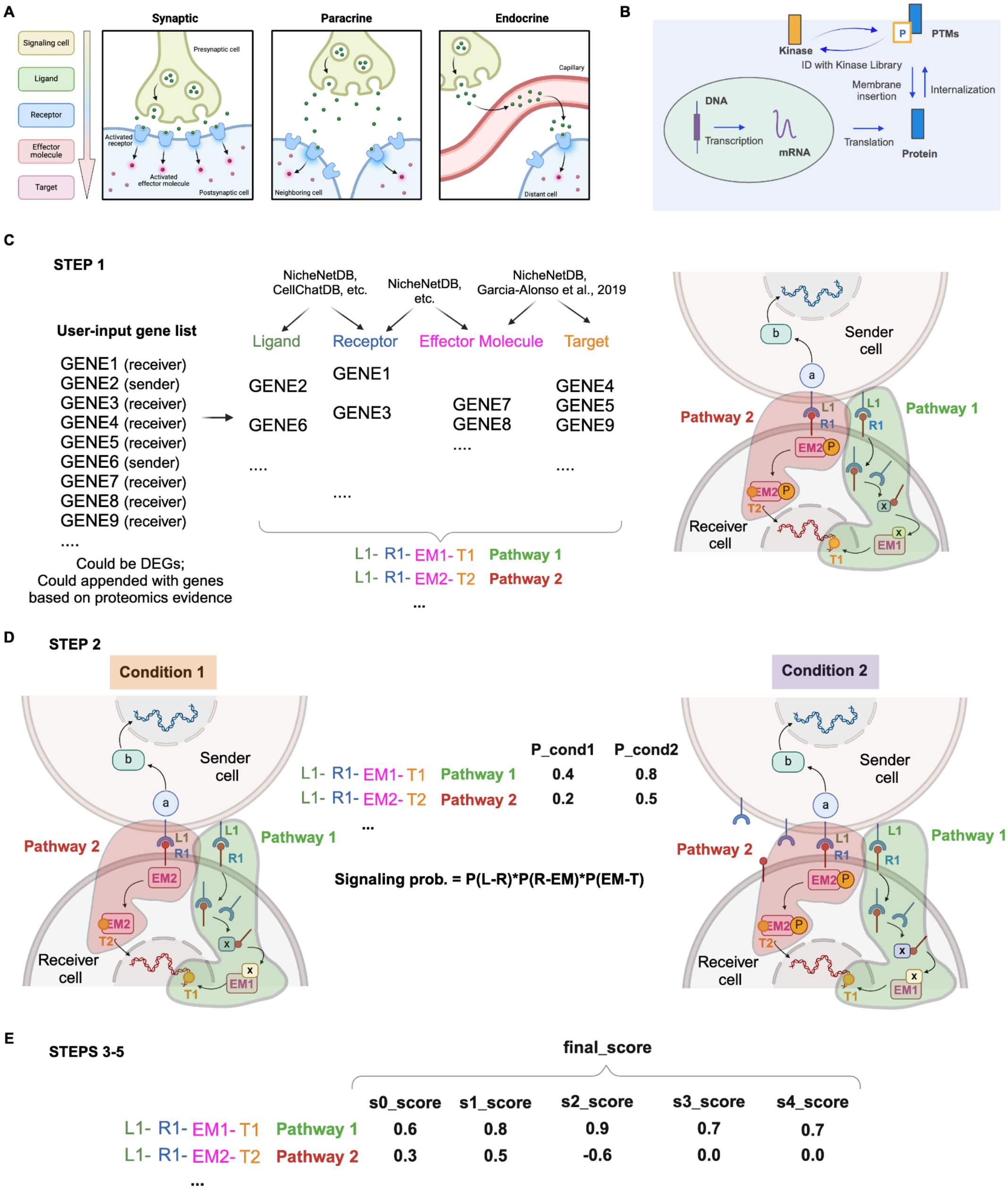


**Supplementary Figure 1. A cartoon illustration depicting the steps of the Incytr algorithm. A.** Since L-R-EM-T pathways are constructed using RNA expression data, the inference captures not only synaptic cell-cell communication but also accommodates paracrine and endocrine interactions. **B.** Incytr's cell signaling pathway inference is supported by evidence gathered from transcription, translation, and protein phosphorylation processes. **C-E.** Incytr pathway inference steps.

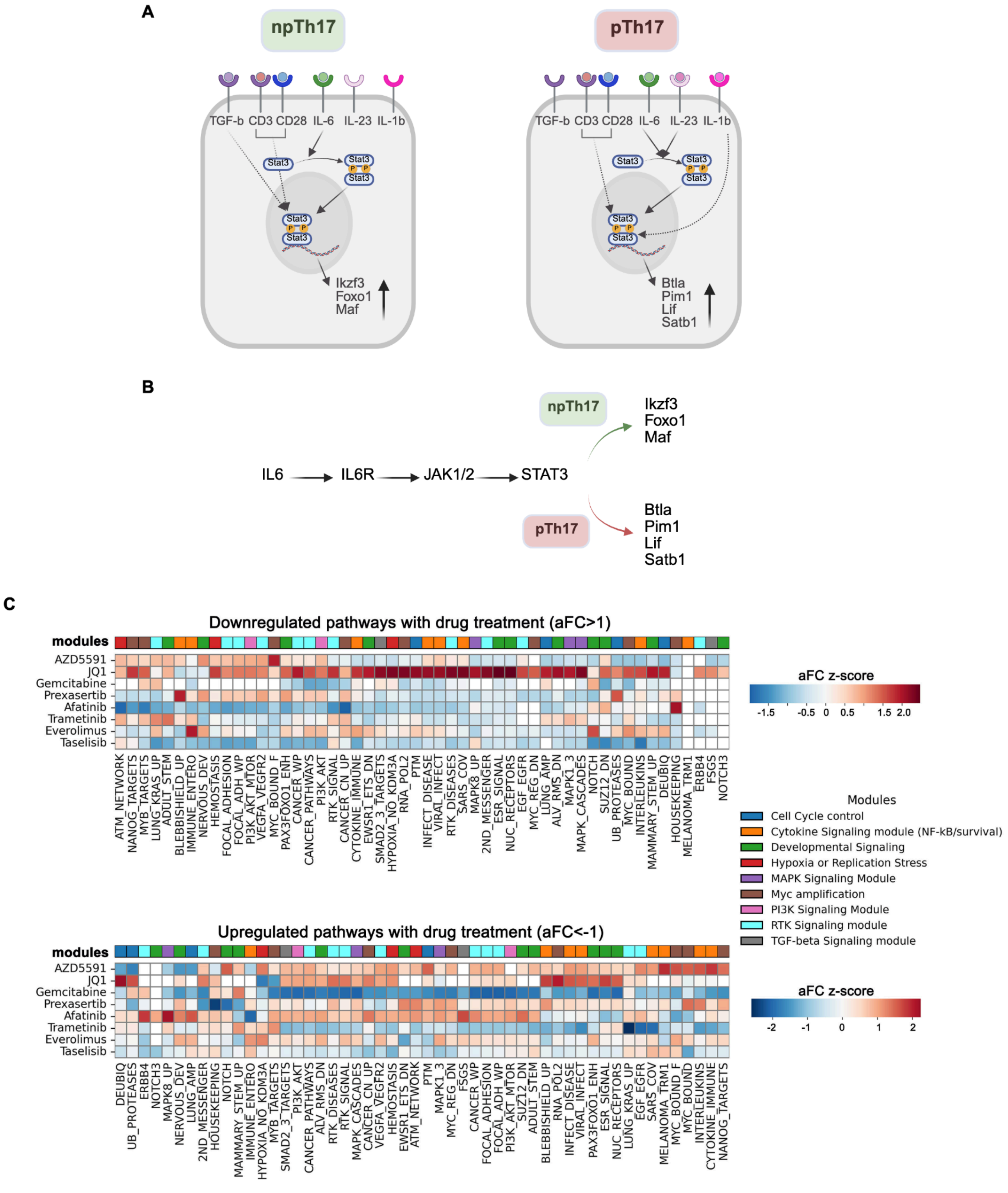
A
npTh17
pTh17
TGF-b CD3 CD28 IL-6 IL-23 IL-1b
Stat3
P
Ikzf3
Foxo1
Maf
Btla
Pim1
Lif
Satb1
B
IL6
IL6R
JAK1/2
STAT3
npTh17
Ikzf3
Foxo1
Maf
pTh17
Btla
Pim1
Lif
Satb1
C
Downregulated pathways with drug treatment (aFC>1)
modules
AZD5591
JQ1
Gemcitabine
Prexasertib
Afatinib
Trametinib
Everolimus
Taselisib
ATM_NETWORK
NANOG_TARGETS
MYB_TARGETS
LUNG_KRAS_UP
ADULT_STEM
BLEBBISHIELD_UP
IMMUNE_ENTERO
NERVOUS_DEV
HEMOSTASIS
FOCAL_ADHESION
FOCAL_ADH_WP
PI3K_AKT_MTOR
VEGFA_VEGFR2
MYC_BOUND_F
PAX3FOXO1_ENH
CANCER_WP
CANCER_PATHWAYS
PI3K_AKT
RTK_SIGNAL
CANCER_CN_UP
CYTOKINE_IMMUNE
EWSR1_ETS_DN
SMAD2_3_TARGETS
HYPOXIA_NO_KDM3A
RNA_POL2
PTM
INFECT_DISEASE
VIRAL_INFECT
RTK_DISEASES
SARS_COV
MAPK8_UP
2ND_MESSENGER
ESR_SIGNAL
NUC_RECEPTORS
EGF_EGFR
MYC_REG_DN
LUNG_AMP
ALV_RMS_DN
MAPK1_3
MAPK_CASCADES
NOTCH
SUZ12_DN
UB_PROTEASES
MYC_BOUND
INTERLEUKINS
MAMMARY_STEM_UP
DEUBIQ
HOUSEKEEPING
MELANOMA_TRM1
ERBB4
FSGS
NOTCH3
aFC z-score
-1.5 -0.5 0 0.5 1.5 2.0
Modules
Cell Cycle control
Cytokine Signaling module (NF-kB/survival)
Developmental Signaling
Hypoxia or Replication Stress
MAPK Signaling Module
Myc amplification
PI3K Signaling Module
RTK Signaling module
TGF-beta Signaling module
Upregulated pathways with drug treatment (aFC<-1)
modules
AZD5591
JQ1
Gemcitabine
Prexasertib
Afatinib
Trametinib
Everolimus
Taselisib
DEUBIQ
UB_PROTEASES
ERBB4
NOTCH3
MAPK8_UP
NERVOUS_DEV
LUNG_AMP
2ND_MESSENGER
HOUSEKEEPING
NOTCH
MAMMARY_STEM_UP
IMMUNE_ENTERO
HYPOXIA_NO_KDM3A
MYB_TARGETS
SMAD2_3_TARGETS
PI3K_AKT
CANCER_PATHWAYS
ALV_RMS_DN
RTK_DISEASES
RTK_SIGNAL
MAPK_CASCADES
CANCER_CN_UP
VEGFA_VEGFR2
HEMOSTASIS
EWSR1_ETS_DN
ATM_NETWORK
PTM
MAPK1_3
MYC_REG_DN
FSGS
CANCER_WP
FOCAL_ADHESION
FOCAL_ADH_WP
PI3K_AKT_MTOR
SUZ12_DN
ADULT_STEM
BLEBBISHIELD_UP
RNA_POL2
INFECT_DISEASE
VIRAL_INFECT
PAX3FOXO1_ENH
ESR_SIGNAL
NUC_RECEPTORS
LUNG_KRAS_UP
EGF_EGFR
SARS_COV
MELANOMA_TRM1
MYC_BOUND_F
MYC_BOUND
INTERLEUKINS
CYTOKINE_IMMUNE
NANOG_TARGETS
aFC z-score
-2 -1 0 1 2

**Supplementary Figure 2. Validation of Incytr in discovering functionally meaningful pathways. A.** Graphical representation of non-pathogenic (npTh17; anti-CD3 + anti-CD28 + IL-6 + TGFβ) and pathogenic (pTh17; anti-CD3 + anti-CD28 + IL-6 + IL-1β + IL-23) stimulation conditions applied to naïve CD4+ T cells, as described [Ariss et al., 2024], and the role of Stat3 in differentiation. **B.** Signaling pathways identified by Incytr 24 hours post-stimulation. Artificial cell cluster X was added to the scRNAseq data with two cells per condition, expressing exclusively Il6 and Il6ra (at an expression level of the maximum expression of any gene in the real data). Incytr was run with X as the sender group. **C.** Heatmap of z-score clustered adjusted fold-change (aFC) from Incytr analysis of scRNAseq from A549 cells treated with inhibitors for 24 h [McFarland et al., 2020]. aFC > 1 denotes downregulated pathways, whereas aFC < –1 denotes upregulated pathways. Pathways are annotated by signaling modules.

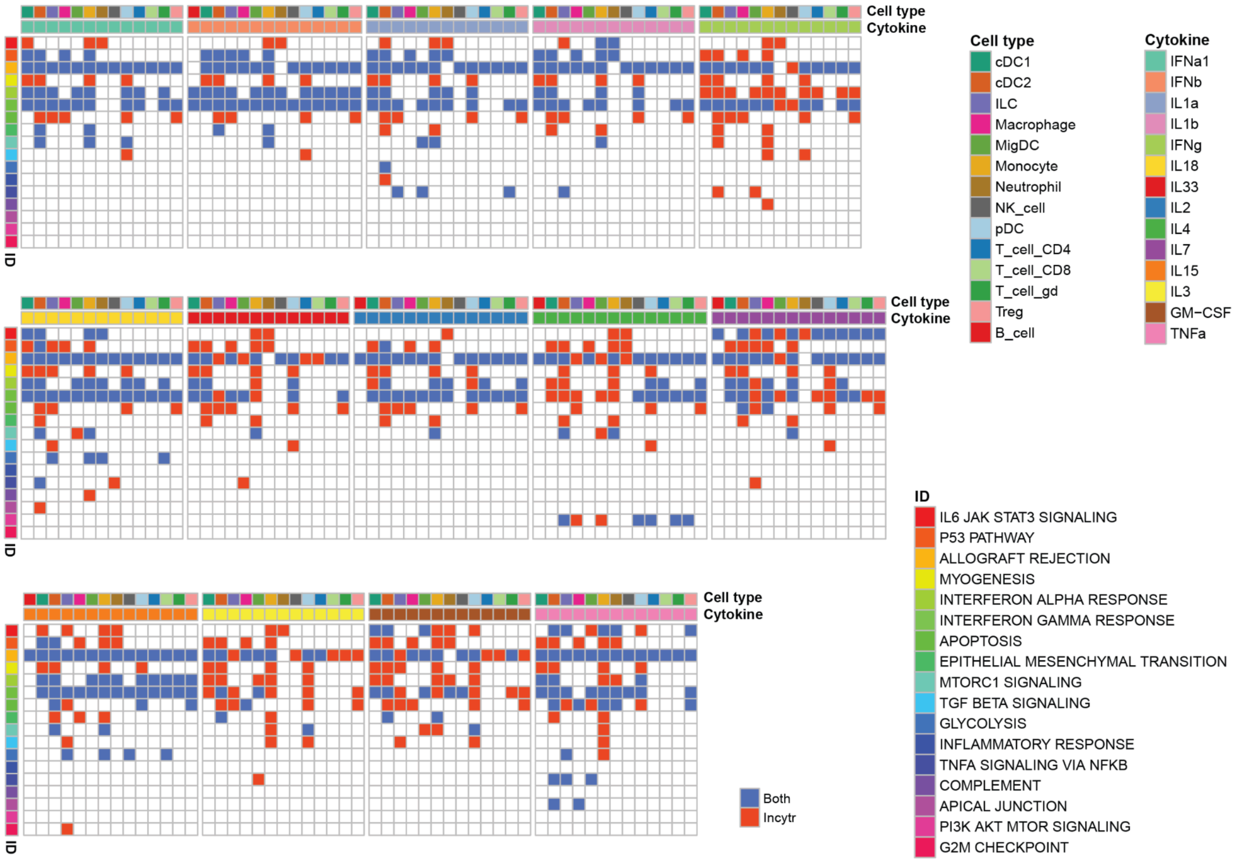


**Supplementary Figure 3. Hallmark IDs identified by Incytr in response to specific cytokine stimulations.** We present results for L → R → EM → T networks that reflect signaling pathways either directly driven by the cytokines (i.e., ligands) used for stimulation or indirectly influenced by them. Incytr-derived networks indirectly influenced by cytokines were restricted to those containing at least one gene reported in Supplementary Table 5 of [Cui et al., 2024]. In the plot, blue indicates Hallmark IDs identified by both Incytr and the cytokine dictionary study, while red denotes those identified by Incytr alone.

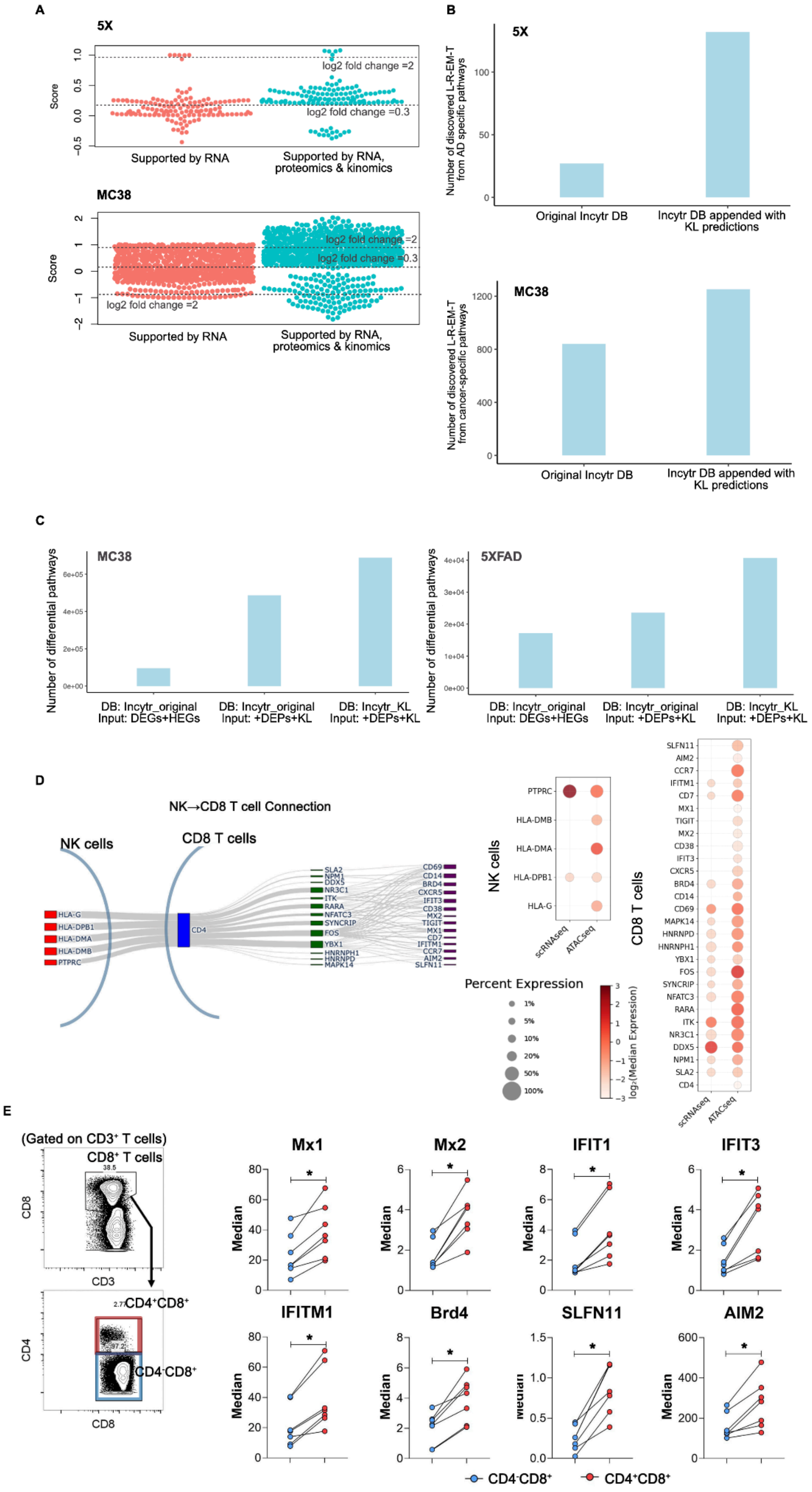
A
5X
Score
log2 fold change =2
log2 fold change =0.3
Supported by RNA
Supported by RNA, proteomics & kinomics
MC38
Score
log2 fold change =2
log2 fold change =0.3
log2 fold change =2
Supported by RNA
Supported by RNA, proteomics & kinomics
B
5X
Number of discovered L-R-EM-T from AD specific pathways
Original Incytr DB
Incytr DB appended with KL predictions
MC38
Number of discovered L-R-EM-T from cancer-specific pathways
Original Incytr DB
Incytr DB appended with KL predictions
C
MC38
Number of differential pathways
DB: Incytr_original Input: DEGs+HEGs
DB: Incytr_original Input: +DEPs+KL
DB: Incytr_KL Input: +DEPs+KL
5XFAD
Number of differential pathways
DB: Incytr_original Input: DEGs+HEGs
DB: Incytr_original Input: +DEPs+KL
DB: Incytr_KL Input: +DEPs+KL
D
NK→CD8 T cell Connection
NK cells
CD8 T cells
HLA-G
HLA-DPB1
HLA-DMA
HLA-DMB
PTPRC
CD4
NK cells
CD8 T cells
scRNAseq
ATACseq
Percent Expression
1%
5%
10%
20%
50%
100%
log₂(Median Expression)
SLFN11
AIM2
CCR7
IFITM1
CD7
MX1
TIGIT
MX2
CD38
IFIT3
CXCR5
BRD4
CD14
CD69
MAPK14
HNRNPD
HNRNPH1
YBX1
FOS
SYNCRIP
NFATC3
RARA
ITK
NR3C1
DDX5
NPM1
SLA2
CD4
E
(Gated on CD3⁺ T cells)
CD8⁺ T cells
CD8
CD3
CD4
CD8
CD4⁺CD8⁺
CD4⁻CD8⁺
Mx1
Mx2
IFIT1
IFIT3
IFITM1
Brd4
SLFN11
AIM2
Median
CD4⁻CD8⁺
CD4⁺CD8⁺

**Supplementary Figure 4. Proteomics and kinomics data enable the reliable rediscovery of known pathways in Alzheimer's disease and cancer. A.** Proteomics and kinomics data provide additional evidence, improving the scoring of pathways that might otherwise (based solely on mRNA data) be considered non-differential between WT and 5XFAD mice in the 5XFAD mouse model, and non-significant in the MC38 mouse cancer model. Each point represents an L-R-EM-T pathway mapped to Alzheimer's disease (AD) or cancer-specific pathways (Focal Adhesion, VEGFR, Cancer Pathways) on WikiPathways. **B.** Appending the Incytr database with PPI predictions from the Kinase Library (KL) enables the discovery of over four times more L-R-EM-T pathways mapped to AD-specific pathways and over one-and-a-half times more L-R-EM-T pathways mapped to cancer-specific pathways on WikiPathways. This approach strengthens the evidence for the presence of these pathways in the given biological system. **C.** The added value of supplementing the input gene list with proteomics evidence (i.e., genes that are differentially expressed at the protein level between condition groups) and appending the Incytr DB (Incytr_original) with the Kinase Library predictions (Incytr_KL). This approach facilitates the discovery of more differential pathways (signaling probability in at least one condition > 0.1, p value in at least one condition < 0.05, and absolute value of the PDS > 0.76). Here, DEGs refer to differentially expressed genes, HEGs to highly expressed genes, DEPs refer to proteins that are differentially expressed between condition groups, and KL refers to the Kinase Library (kinase-substrate predictions). **D.** Incytr analysis of ATACseq data from a cohort of individuals living with HIV identifies NK→CD8 T cell signaling via MHC-II-CD4 ligand-receptor interactions across a four-step network, with downstream target gene activation. **E.** Orthogonal CyTOF validation in the same cohort showing paired, median protein expression of selected markers in $CD4^{-}CD8^{+}$ T cells (blue) versus $CD4^{+}CD8^{+}$ T cells (red). * , *p*-value <0.05 (non-paramteric Wilcoxon test).

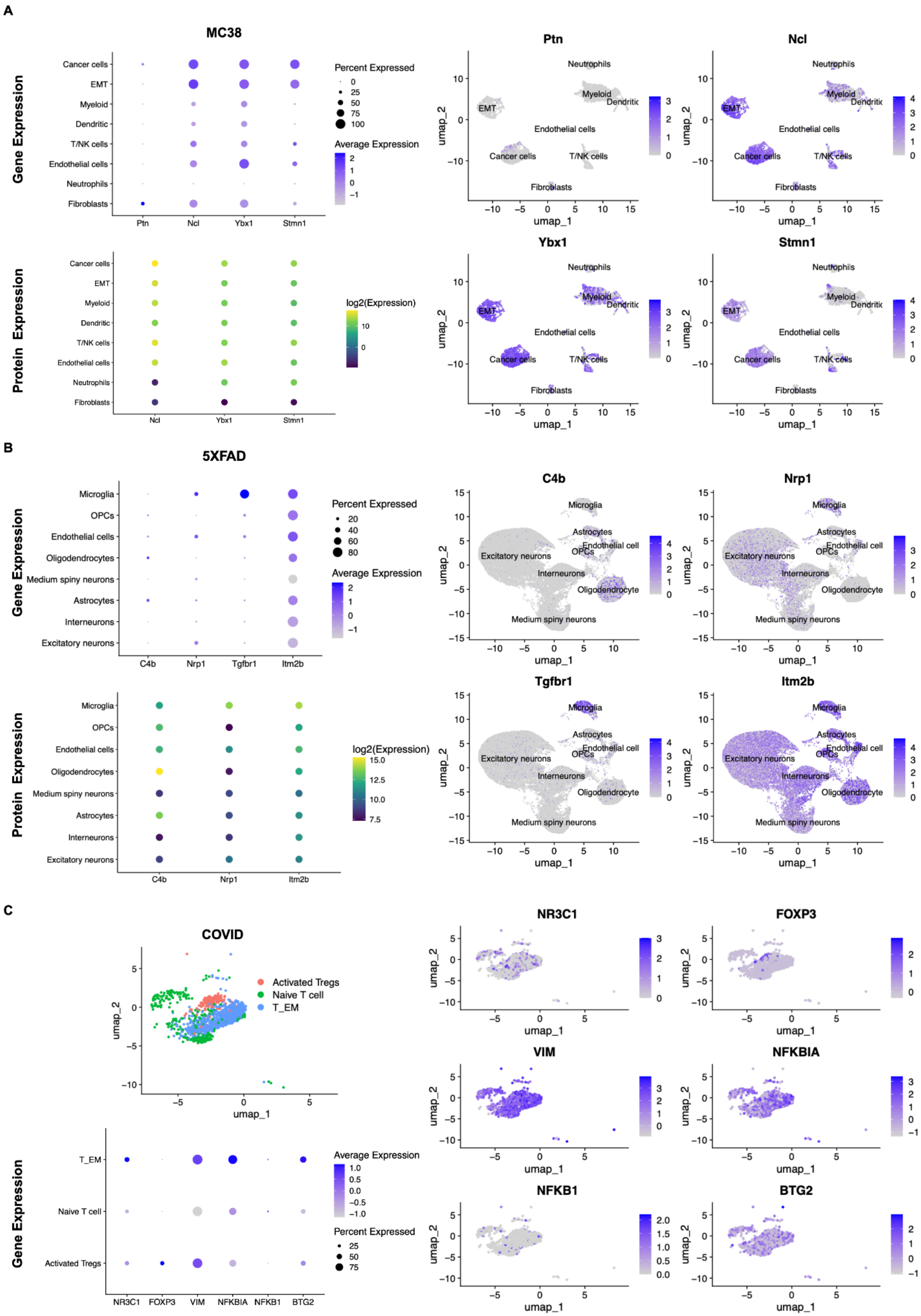
A
MC38
Gene Expression
Protein Expression
Cancer cells
EMT
Myeloid
Dendritic
T/NK cells
Endothelial cells
Neutrophils
Fibroblasts
Ptn
Ncl
Ybx1
Stmn1
Percent Expressed
Average Expression
log2(Expression)
umap_1
umap_2
B
5XFAD
Microglia
OPCs
Endothelial cells
Oligodendrocytes
Medium spiny neurons
Astrocytes
Interneurons
Excitatory neurons
C4b
Nrp1
Tgfbr1
Itm2b
Endothelial cell
Oligodendrocyte
C
COVID
Activated Tregs
Naive T cell
T_EM
NR3C1
FOXP3
VIM
NFKBIA
NFKB1
BTG2

**Supplementary Figure 5. Gene and protein expression patterns for representative Incytr-inferred signaling pathways in the MC38 (A), 5XFAD (B), and primary patient cohort in COVID-19 (C) datasets.**

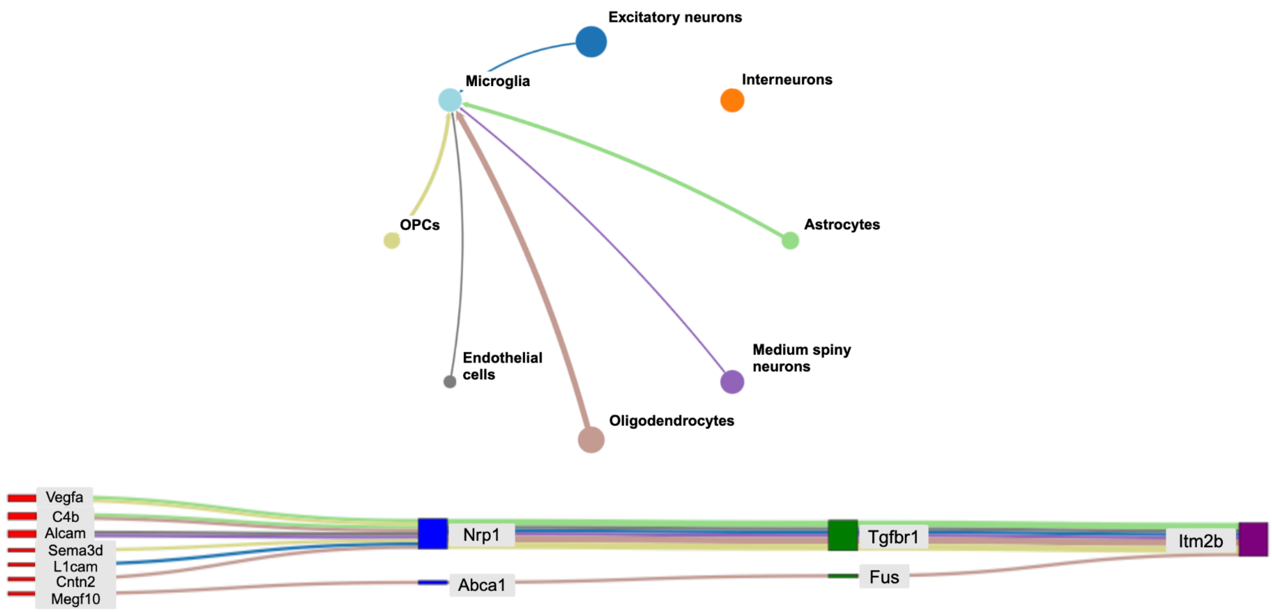


**Supplementary Figure 6. Incytr-identified pathways showing differential expression (log2 fold change > 0.5 for T-PDS and PDS scores) in 5XFAD versus WT mice, involving the Itm2b gene in microglial cells.**

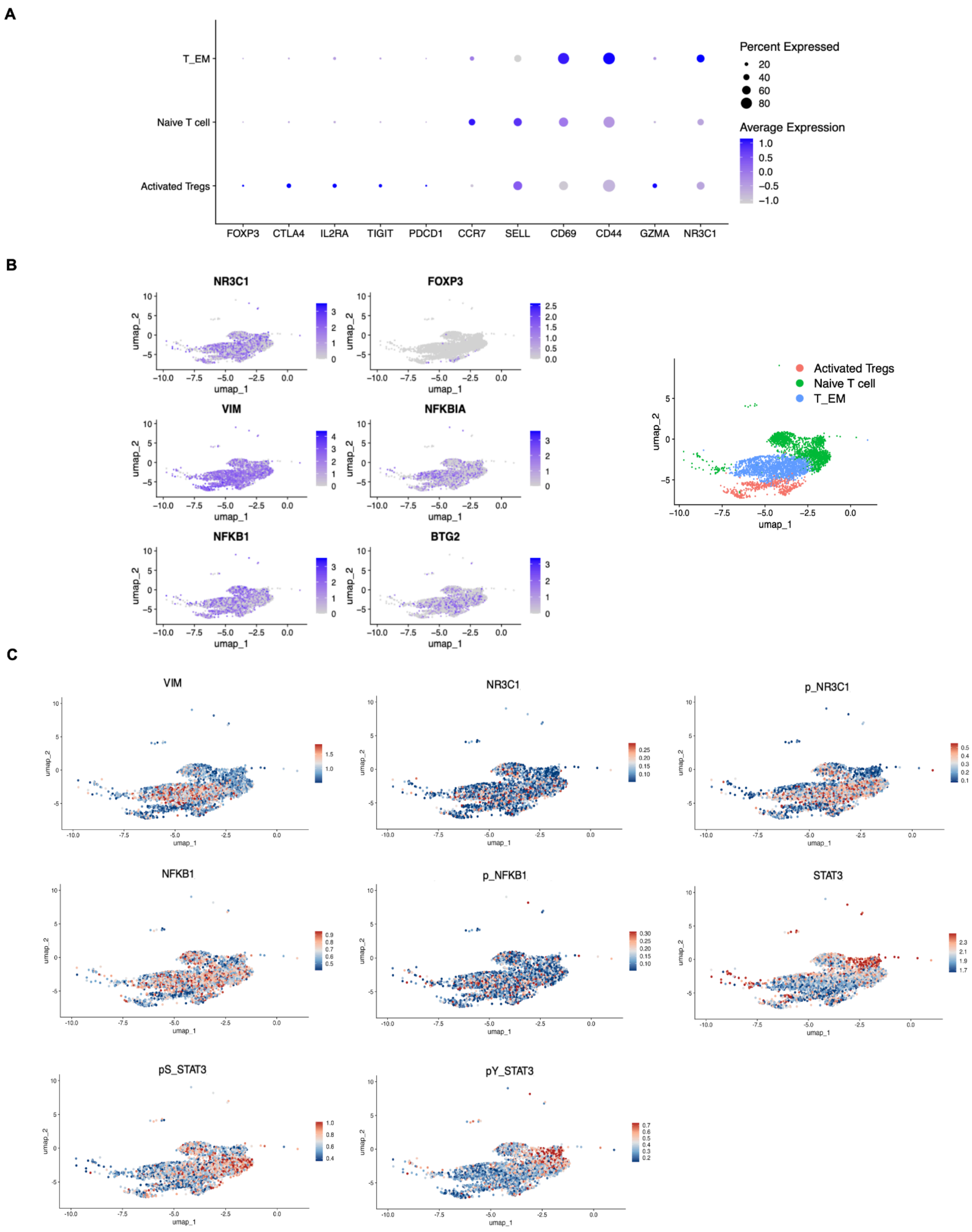


**Supplementary Figure 7. Protein expression patterns in the second cohort of COVID-19 patients. A.** Subclustering of the T cell population in this cohort aligns with T cell phenotyping from the first cohort shown in Figure 5B. **B.** mRNA expression profiles of the selected genes. **C.** InTraSeq results for selected proteins.

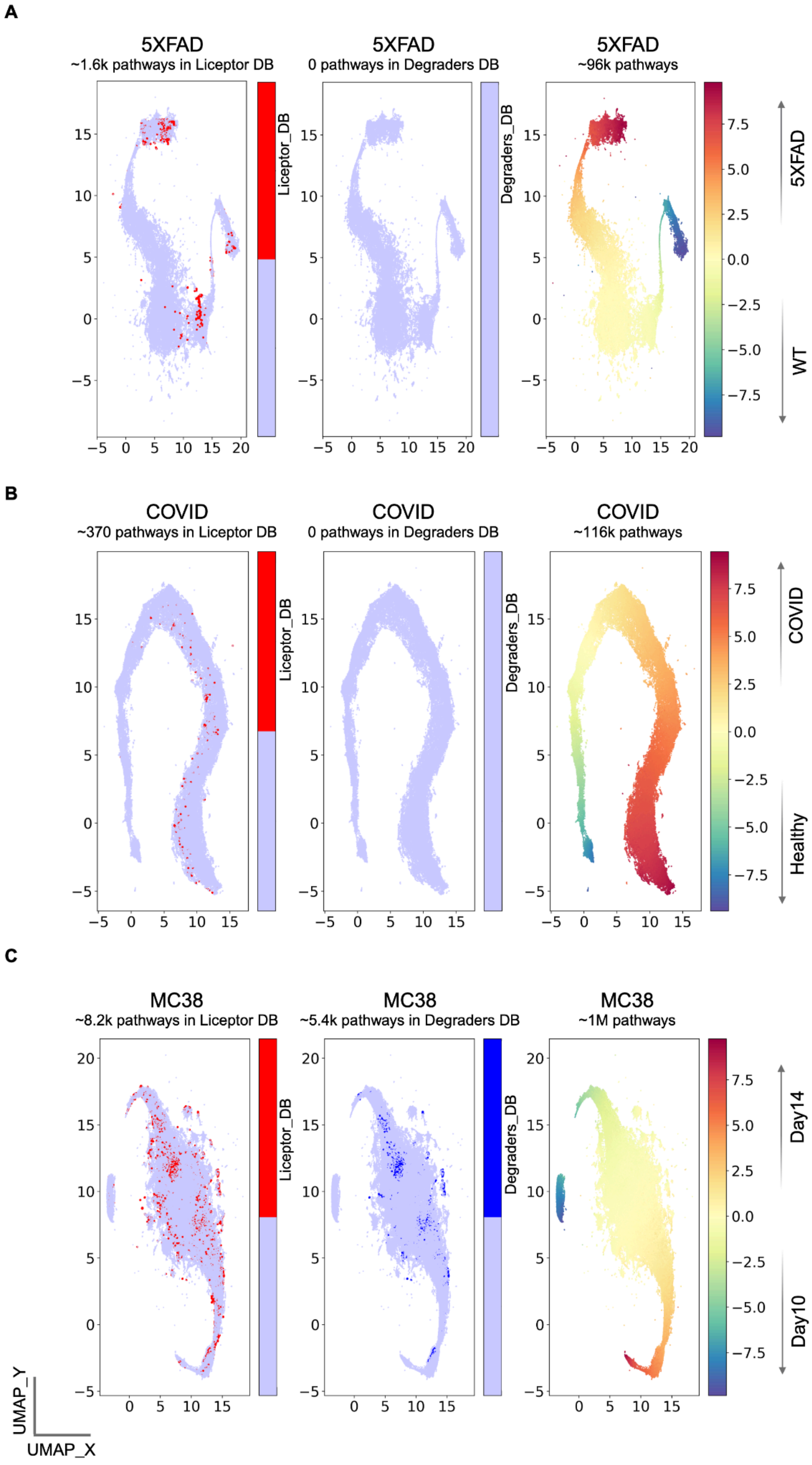
A
5XFAD
~1.6k pathways in Liceptor DB
Liceptor_DB
5XFAD
0 pathways in Degraders DB
Degraders_DB
5XFAD
~96k pathways
5XFAD
WT
B
COVID
~370 pathways in Liceptor DB
Liceptor_DB
COVID
0 pathways in Degraders DB
Degraders_DB
COVID
~116k pathways
COVID
Healthy
C
MC38
~8.2k pathways in Liceptor DB
Liceptor_DB
MC38
~5.4k pathways in Degraders DB
Degraders_DB
MC38
~1M pathways
Day14
Day10
UMAP_Y
UMAP_X

**Supplementary Figure 8. The number of L-R-EM-T pathways annotated in the Liceptor and Degraders databases.** Signaling pathways containing at least one molecule (L, R, EM, or T) annotated in the Liceptor or Degraders databases are highlighted in red or blue, respectively, in the 5XFAD (**A**), COVID (**B**), and MC38 (**C**) datasets.

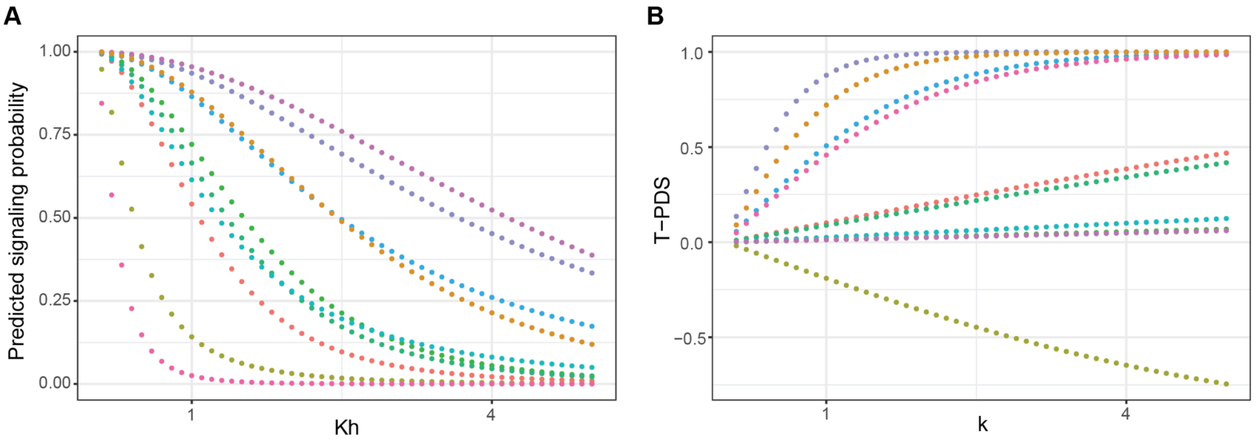


**Supplementary Figure 9. The parameter perturbations of the signaling probability prediction and T-PDS calculation, respectively. A.** Predicted signaling probabilities with different $Kh$ values, showing that changing the value of $Kh$ from 0.1 to 5 (with step size 0.1) does not change the order of signaling probabilities of the L-T pathways. Dot plot showing the predicted signaling probabilities of the synthetic data, where each gene has a random expression value between 0.1 to 5. **B.** Calculated T-PDS with different *k* values, showing that changing the value of *k* from 0.1 to 5 (with step size 0.1) does not change the order of T-PDS of the L-T pathways. Dot plot showing the calculated T-PDS of the synthetic data, where the signaling probabilities of each L-T pathway in two conditions are randomly picked between 0.1 and 1.

| Concept | Definition | References |
|---|---|---|
| effector molecule (EM) | intermediate signal transmission molecule | New |
| target | molecule/gene that is regulated by the effector molecule | [Browaeys et al., 2019] |
| ligand-target signaling pathway | signaling pathway that starts from the ligand to the target molecule/gene | [Browaeys et al., 2019] |
| ligand-target signaling pathway (L-T pathway) | pathway that has the “ligand-receptor-EM-target” structure | New |
| signaling-involved kinase (SiK) | kinase which itself and its substrate are both the receptor/EM/target of an L-T pathway | New |
| signaling-related kinase (SrK) | kinase which is not SiK but predicted by KL to modify the substrate (receptor/EM/target of an L-T pathway) | New |
| adjusted fold-change value (aFC value) | adjusted log2 fold change to turn down the significant change made by two small numbers | New |
| exclusiveness index (EI) | the index that quantifies how exclusively a gene is expressed in the cell group | New |
| SiK-score | the score that quantifies how much a SiK contributes to the L-T pathway expression | New |
| transcriptomics-based pathway differential score (T-PDS) | the score is based on the analysis using the transcriptomics data and quantifies how an L-T pathway is differentially expressed between conditions | New |
| proteomics-based pathway differential score (P-PDS) | the score is based on the analysis using the proteomics data and quantifies how an L-T pathway is differentially expressed between conditions | New |
| phosphorylation-based pathway differential score (Ph-PDS) | the score is based on the analysis using the phosphorylation data and quantifies how an L-T pathway is differentially expressed between conditions | New |
| multi-modal differential | the score that quantifies how an L-T pathway | New |

| | | |
|---|---|---|
| score | is differentially expressed between conditions based on the T-PDS, P-PDS, and Ph-PDS | |
| kinase-based pathway differential score (K-PDS) | the score that quantifies how much the identified SiKs contribute to the L-T pathway expression | New |
| pathway differential score (PDS) | the score that quantifies how an L-T pathway is differential between the conditions (includes T-PDS, P-PDS, Ph-PDS, and K-PDS) | New |

**Supplementary Table 1. A list of terminologies and concepts used in Incytr.**

| | Ligand-Receptor | Receptor-EM | EM-Target |
|---|---|---|---|
| Human | 6869 | 3757666 | 4678245 |
| Mouse | 6707 | 3717790 | 4441402 |

**Supplementary Table 2. The number of interactions recorded in the Incytr-DB.**

| Elemental Isotope | Antigen Target | Vendor | Catalog Number |
|---|---|---|---|
| 115In | CD14 | Biolegend | 301843 |
| 142Nd | CD8 | Biolegend | 344727 |
| 145Nd | IFITM1* | Proteintech | 60074-1-Ig |
| 146Nd | IFIT1* | Novus | NBP2-71005 |
| 147Sm | CD7 | Santa CruzBiotech | 3147006B |
| 149Sm | AIM2* | Santa Cruz Biotech | sc-293174 |
| 150Nd | IFIT3* | Novus | NBP2-71006 |
| 151Eu | SLFN11* | Cell Signaling Tech | 34858 |
| 153Eu | MX2* | Santa CruzBiotech | sc-271527 |
| 155Gd | pIRF3* | Cell Signaling Tech | 29047 |
| 156Gd | MX1* | Cell Signaling Tech | 62815 |
| 170Er | CD3 | Standard Biotools | 3170001B |
| 173Yb | BRD4* | Abcam | ab182446 |
| 174Yb | CD4 | Standard Biotools | 3174004B |
| 176Yb | CD56 | Standard Biotools | 3176008B |
| 209Bi | CD16 | Standard Biotools | 3209002B |
| *Intracellular | | | |

**Supplementary Table 3. CyTOF Antibodies.**

**Supplementary Data 1.** Benchmarking table comparing the ability of GSEA, exFINDER, and Incytr to rediscover known cancer-specific, 5XFAD model-specific, and COVID-specific pathways.

**Supplementary Data 2.** A list of references supporting the primary pairwise L-R, R-EM, and EM-T interactions presented in this paper (Figures 3-5).

**Supplementary Data 3.** Output of the Incytr analysis on the 5XFAD dataset, focusing on interactions involving Apoe, App, Trem2, and Lrp1.
